# Upstream open reading frames dynamically modulate CLOCK protein translation to regulate circadian rhythm and sleep

**DOI:** 10.1101/2023.11.22.568359

**Authors:** Yuanqiang Sun, Ke Shui, Qinyu Li, Chenlu Liu, Wanting Jin, Jian-Quan Ni, Jian Lu, Luoying Zhang

## Abstract

The circadian rhythm is an evolutionarily conserved mechanism with translational regulation increasingly recognized as pivotal in its modulation. In this study, we found that upstream open reading frames (uORFs) are enriched in circadian rhythm genes, with particularly conserved uORFs present in core circadian clock genes. We demonstrate evidence that the uORFs of the core clock gene, *Clock* (*Clk*), rhythmically and substantially attenuate CLK protein translation in *Drosophila*, with pronounced suppression occurring during daylight hours. Eliminating *Clk* uORFs results in elevated CLK protein levels during the day and a compressed circadian cycle, along with a broad shift in clock gene expression rhythms. Interestingly, *Clk* uORF deletion also augments morning sleep by reducing dopaminergic activity. Beyond daily circadian adjustments, *Clk* uORFs play a role in modulating sleep patterns in response to the varying day lengths of different seasons, inhibiting translation in a day-length contingent manner. Furthermore, the *Clk* uORFs act as a master regulator to shape the rhythmic expression of a vast array of genes and influence multifaceted physiological outcomes. Collectively, our research sheds light on the intricate ways uORFs dynamically adjust downstream coding sequences to acclimate to environmental shifts.

## Introduction

Circadian rhythms orchestrate ∼24-hour cycles in a multitude of biological processes, with the sleep-wake cycle being a particularly salient manifestation. The molecular underpinnings of the circadian rhythms consist of a transcription-translation feedback loop (TTFL), an evolutionarily conserved mechanism evident across diverse taxa, ranging from bacteria to fungi, plants, and animals ^1–4^. The circadian systems drive gene expression rhythms of thousands of genes, exerting pervasive influences on daily fundamental biological activities from the cellular level to complex behaviors and diseases such as cell proliferation, cell signaling, metabolism, rest/activity, cancer and neurodegeneration ^5–7^. Despite shared rhythmic principles in the tree of life, species-specific divergence in clock components and mechanisms is evident, likely a result of distinct environmental pressures and life histories shaping unique adaptations after a single origin in a common ancestor of all life forms ^4,8^. Alternatively, the difference in genetic components across different kingdoms of life implies that these rhythmic systems probably emerged separately in various lineages at an early stage via convergent evolution ^9,10^. While the key genetic components that drive the circadian rhythms have been deciphered ^5–7^, the intricate regulatory mechanisms fine-tuning these rhythms remain partially obscured, with significant gaps in our understanding persisting.

*Drosophila*, with its genetic tractability and conservation of key circadian components, serves as an unparalleled model providing profound insights that resonate across species, including humans ^4,11–14^. In *Drosophila*, two transcriptional activators, CLOCK (CLK) and CYCLE (CYC), constitute the positive limb of the loops. CLK and CYC heterodimerize and activate the transcription of *period* (*per*) and *timeless* (*tim*) via E-box enhancer elements ^15,16^. As PER and TIM proteins accumulate in the cytoplasm, they bind to each other and translocate into the nucleus to suppress the transcriptional activities of CLK and CYC, thus repressing their own transcription and forming a pivotal negative feedback loop ^17^. Throughout this process, the stoichiometry of these core regulators is precisely controlled to ensure the accuracy of the circadian rhythms. For instance, the overexpression of CLK-CYC increases their activity at *per* and *tim* E-box enhancers, consequently shortening the circadian rhythms ^18,19^. Recent studies have revealed that the precision of circadian rhythms is regulated by multi-layered mechanisms beyond transcriptional modulation, which include the regulation of protein synthesis and post-translational stability. For instance, the translation of *tim* and *per* mRNAs is facilitated by TWENTY-FOUR and its activator, ATAXIN2, through their promotion of an association with polyadenylate-binding protein ^20–23^. Additionally, the synthesis of CLK and TIM proteins is inhibited by the microRNAs *bantam* ^24^ and *mir-276a* ^25^, respectively, with the overexpression of either miRNA leading to prolonged periods or arrhythmicity ^24–26^. Post-translational modifications (PTMs) also affect the stability and subsequent degradation of PER and TIM ^1^.

While recent advances have significantly enhanced our understanding of the regulatory complexity of the classical TTFL, the field has not yet completely elucidated the intricate network of components and interactions that govern the fine-scale regulation of circadian rhythms, even in *Drosophila*, the quintessential model for circadian biology.

Recent studies underscore the pivotal role of translational regulation, particularly by upstream open reading frames (uORFs), in the control of circadian rhythms in organisms with divergent circadian architectures. uORFs, the short ORFs with start codons located within the 5’ untranslated regions (5’UTRs) of eukaryotic mRNAs, are widely distributed in eukaryotic genomes ^27^. uORFs play a key role in suppressing translation initiation of the downstream coding sequence (CDS) in an mRNA by either sequestering ribosomes or competing for binding to translation initiation complexes ^28–32^. In *Neurospora*, the central circadian clock component, *frequency* (*frq*), has two isoforms—large (l-FRQ) and small (s-FRQ)—whose balance is determined by temperature-dependent splicing and is essential for the temperature compensation of circadian rhythms. Thermosensitive trapping of scanning ribosomes at the uORFs of l-FRQ modulates FRQ protein levels in response to ambient temperatures, thus calibrating the circadian clock ^33^. Similarly, TIMING OF CAB EXPRESSION 1 (TOC1), a core component of the *Arabidopsis* circadian clock, is regulated by uORFs within its mRNA leader sequence ^34^. Although *Neurospora* and *Arabidopsis* share similar design principles in circadian rhythms as humans and flies, the proteins involved differ significantly in both structure and conservation ^35–37^. Recent studies have shown that uORFs in the mouse core clock gene *Period2* (*Per2*) play a role in temperature entrainment of the circadian clock ^38^ and in regulating sleep ^39^. However, no significant differences in the period and amplitude of circadian rhythms were observed between wild-type and *Per2* uORF mutant mice ^39^. In addition, extensive ribosomal binding to uORFs in rhythmically expressed mRNAs has been observed in mammalian cells, yet the functional impacts of these uORFs in circadian regulation have not been characterized ^40,41^. In summary, although research into the translational regulation by uORFs is burgeoning, the complete spectrum of regulatory functions of uORFs in circadian rhythm control remains to be elucidated. Moreover, the potential impacts of uORF-mediated regulation on physiological and pathological processes linked to circadian rhythms are currently elusive, while their delineations can help reveal how circadian rhythms adapt to environmental pressures.

Here, we show that circadian rhythm-related genes are significantly enriched with ultra-conserved uORFs in *Drosophila*. These uORFs, which demonstrate translational activity in our previously published translatome data ^42^, thus may play a crucial role in modulating circadian rhythms. We then conducted an in-depth exploration of conserved and potentially functional uORFs in *Clk* of *Drosophila melanogaster* and presented evidence that these uORFs substantially inhibit protein translation *in vitro* and *in vivo*. Knocking out (KO) *Clk* uORFs amplifies daytime CLK protein levels and shortens the circadian period of locomotor rhythm. Furthermore, *Clk-*uORF-KO mutants manifest increased morning sleep, likely stemming from diminished dopaminergic activity. These mutants also exhibit defects in modulating sleep duration in response to photoperiodic shifts. Molecular investigations underscore the requirement of *Clk* uORFs for photoperiod-driven regulation of CLK protein, highlighting their central role in both circadian and circannual orchestrations. Intriguingly, the mutants also display decreased fecundity and diminished resilience to starvation, indicating that the ablation of *Clk* uORFs has extensive physiological consequences beyond the regulation of circadian rhythms and sleep.

## Results

### Circadian rhythm genes are more likely to be regulated by uORFs

We identified 36,565 putative canonical uORFs across all isoforms of 13,470 protein-coding genes in *D. melanogaster*. Of these, 1,137 uORFs are associated with 169 circadian rhythm-related genes (GO:0007623), demonstrating a significant enrichment of uORFs in this category of genes (*P* < 10^-16^, Fisher’s exact test; Fig. 1A). Since conserved uORFs are more likely to be functional, we then focused on uORFs highly conserved across *Drosophila* species. As the start codon (uATG) is the most pivotal and definitive feature of a canonical uORF ^30,43^, we investigated the conservation patterns of the uATGs based on the multiple sequence alignments (MSA) of 23 *Drosophila* species. This analysis revealed 388 uATGs (within 250 genes) that are identical across the *Drosophila* species examined (Table S1). These genes are significantly enriched for functions in neuronal axon genesis and transcriptional regulation, implicating the potential roles of uORFs in modulating these processes (Fig. S1). Among the 169 circadian rhythm-related genes, 29 uORFs display conservation of uATGs, a higher ratio than that of uORFs in the remaining genes (29/1,137 vs. 359/35,428; *P* = 2.3×10^-5^, Fisher’s exact test; Fig. 1B). For the ten core clock genes (Table1), we found that 7 out of 82 uORFs possess uATGs that are identical across *Drosophila* species, suggesting higher uORF conservation in the core versus other circadian genes (7/82 vs. 22/1,055; *P* = 0.005, Fisher’s exact test; Fig. 1B). These seven conserved uORFs are distributed among four genes, with *Pdp1* containing four, and *Clk*, *sgg*, and *cwo* each containing one (Table 1). These patterns suggest that circadian rhythm genes, especially core clock genes, are more inclined to be regulated by uORFs.

**Fig. 1.**
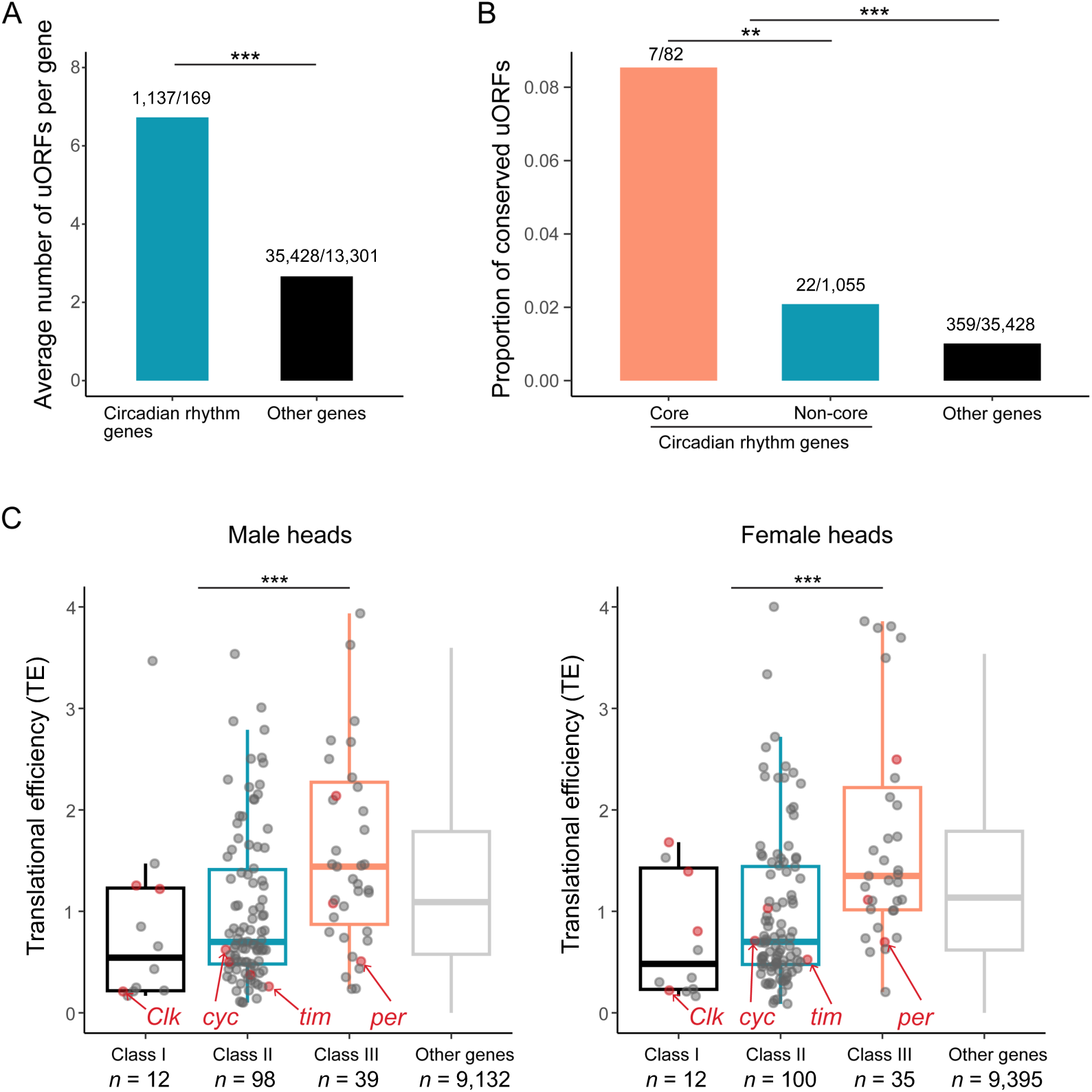
Circadian rhythm genes are more likely to be regulated by uORFs. (A) Average number of uORFs per gene for circadian rhythm-related genes and the other genes in *D. melanogaster*. The ratio of uORFs to genes is denoted on each bar (Fisher’s exact test. ***, *P* < 0.001). (B) Proportion of conserved uORFs in core clock genes, other circadian rhythm genes, and the other genes. Ratios reflect the number of conserved uORFs relative to the total count of uORFs for each category, depicted on each bar (Fisher’s exact test. **, *P* < 0.01; ***, *P* < 0.001). (C) The translational efficiency (TE) for different classes of circadian rhythm-related genes in male and female heads of *D. melanogaster*. Class I includes genes with conserved and translated uORFs. Class II consists of genes with non-conserved yet translated uORFs. Class III comprises genes without translated uORFs or lacking uORFs altogether. ‘Other’ encompasses all remaining genes expressed in heads that are not related to circadian rhythm. The gene counts of each class are denoted at the bottom of each bar. The ten core clock genes are highlighted with red dots, with *Clk*, *cyc*, *tim*, and *per* specifically marked by red arrows (Wilcoxon signed-rank test. ***, *P* < 0.001).

**Table 1.** uORFs in circadian rhythm genes.

| Description | Circadian | Core |  |  |  |  |  |  |  |  |  |  |
| --- | --- | --- | --- | --- | --- | --- | --- | --- | --- | --- | --- | --- |
|  | rhythm | rhythm |  |  |  |  |  |  |  |  |  |  |
|  | genes | genes | *Clk* | *sgg* | *cwo* | *Pdp1* | *tim* | *cyc* | *vri* | *cry* | *Bdbt* | *per* |
| Total uORFs | 1137 | 82 | 7 | 18 | 9 | 24 | 8 | 4 | 12 | 0 | 0 | 0 |
| Conserved uORFs | 29 | 7 | 1 | 1 | 1 | 4 | 0 | 0 | 0 | 0 | 0 | 0 |
| **Male head** |  |  |  |  |  |  |  |  |  |  |  |  |
| uORFs with mRNA RPKM > 0.1 | 888 | 67 | 6 | 15 | 8 | 24 | 6 | 3 | 5 | 0 | 0 | 0 |
| uORFs with TE > 0.1 | 692 | 52 | 5 | 13 | 6 | 15 | 5 | 3 | 5 | 0 | 0 | 0 |
| Conserved uORFs with mRNA RPKM > 0.1 | 28 | 7 | 1 | 1 | 1 | 4 | 0 | 0 | 0 | 0 | 0 | 0 |
| Conserved uORFs with TE > 0.1 | 25 | 7 | 1 | 1 | 1 | 4 | 0 | 0 | 0 | 0 | 0 | 0 |
| **Female head** |  |  |  |  |  |  |  |  |  |  |  |  |
| uORFs with mRNA RPKM > 0.1 | 879 | 63 | 6 | 15 | 8 | 24 | 5 | 3 | 2 | 0 | 0 | 0 |
| uORFs with TE > 0.1 | 714 | 49 | 6 | 13 | 5 | 16 | 5 | 3 | 1 | 0 | 0 | 0 |
| Conserved uORFs with mRNA RPKM > 0.1 | 28 | 6 | 1 | 1 | 0 | 4 | 0 | 0 | 0 | 0 | 0 | 0 |
| Conserved uORFs with TE > 0.1 | 25 | 6 | 1 | 1 | 0 | 4 | 0 | 0 | 0 | 0 | 0 | 0 |

The number of total uORFs or conserved uORFs in 169 circadian rhythm genes and ten core clock genes, as well as each core clock gene, is shown. The number of transcribed and translated uORFs in heads of male and female flies is shown. Conserved uORFs refer to the uATGs of uORFs that are present in all 23 *Drosophila* species. TE, translational efficiency; RPKM, reads per kilobase of transcript per million mapped reads.

Besides conservation, the regulatory functions of uORFs are also implied by their expression and translation patterns. Analysis of our previously published transcriptome and translatome data ^42^ indicates that 78% of the 1,137 uORFs in the circadian rhythm-related genes are expressed in adult heads (mRNA RPKM > 0.1, 888 in males and 879 in females; Table 1), and about 80% of these expressed uORFs display evidence of translation [Translation efficiency (TE) > 0.1, 692/888 in males and 714/879 in females; Table 1]. Notably, 25 of the 29 highly conserved uORFs exhibit translation signals in the heads of both sexes. Of the 82 uORFs from the core clock genes, 60% show mRNA expression in adult heads (67 in males and 63 in females), with 78% of these expressed uORFs exhibiting translation signals (52/67 in males and 49/63 in females; Table 1), suggesting their potential functional importance.

To assess the impact of uORFs on translation efficiency of the downstream CDS, we categorized the circadian rhythm-related genes expressed in heads (149 in male and 147 in female) into three groups: Class I (genes with conserved uORFs that are translated, 12 in male and 12 in female), Class II (genes with translated uORFs that are not highly conserved, 98 in male and 100 in female), and Class III (genes without translated uORFs, 39 in male and 35 in female). We observed that genes with translated uORFs (Class I and II) overall display a lower TE than those without (Class III) (*P* < 0.0002 for both male and female heads, respectively, Wilcoxon signed-rank test; Fig. 1C). While not statistically significant, Class I genes show a tendency of lower TE compared to that of Class II in both sexes. Each of the four core clock genes (*Clk*, *cyc*, *per*, and *tim*) demonstrated reduced TE relative to the genomic average. Notably, *Clk* belongs to Class I and exhibits a significantly lower TE, implying a substantial regulatory impact of uORFs on its translation (Fig. 1C).

### uORFs suppress CLK and CYC translation *in vitro* and *in vivo*

In *Drosophila melanogaster*, the 5’UTR of *Clk* mRNA contains five canonical uORFs, among which the second (uORF2) and the third uORF (uORF3) are in-frame to each other and overlapping (Fig. 2A). Among the five uORFs of *Clk*, the start codon of uORF2 (uATG2) is the most conserved, identical across all 23 *Drosophila* species (Fig. 2A). As CYC is the binding partner of CLK, we also analyzed *cyc* for comparison. The 5’UTR of *cyc* mRNA contains four canonical uORFs (SI Appendix, Fig. S2A). The uATGs of *cyc* are less conserved than those of *Clk* in general, and none of them is identical across the 23 *Drosophila* species (SI Appendix, Fig. S2A). We detected prominent mRNA expression and ribosome occupancy for uORF2 and uORF3 of *Clk* in the head and body of male and female flies (Fig. 2B). As for *cyc*, prominent mRNA expression and ribosome occupancy was observed for uORF2, uORF3 and uORF4 (SI Appendix, Fig. S2B).

**Fig. 2.**
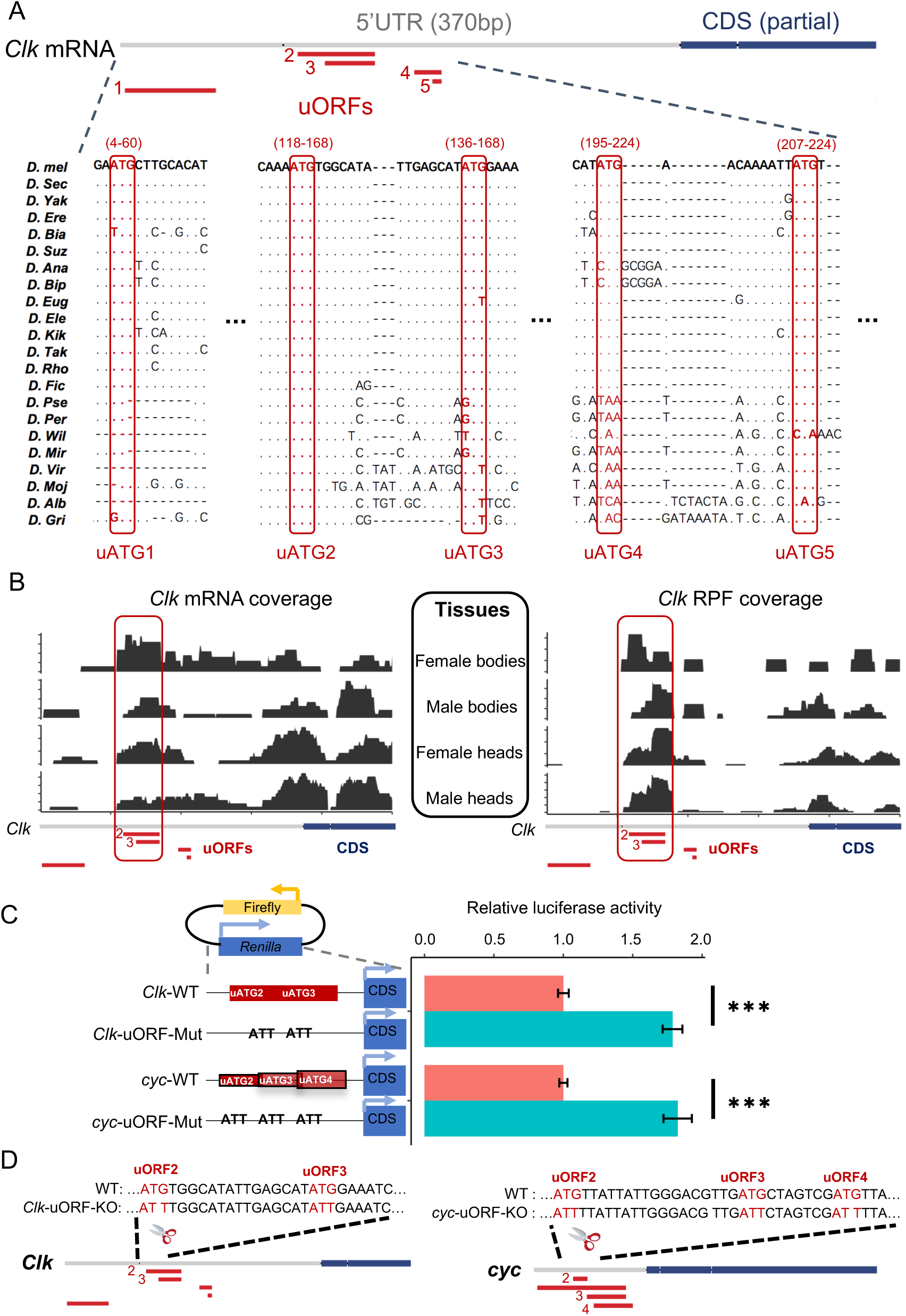
Regulation of CLK and CYC Translation by uORFs in *Drosophila*. (A) Multiple sequence alignment (MSA) of *Clk* uORFs among 23 *Drosophila* species. The start codons (uATGs) of uORFs are highlighted by red boxes. The position schemes of uORFs and partial CDSs are denoted above the MSA, with red and blue colors, respectively. The start and end positions of each uORF (separated by a “-”) in the 5’ UTR are given in the parenthesis above each uATG. (B) The mRNA reads coverage (left) and ribosome-protected footprints (RPF) coverage (right) of uORFs of *Clk* mRNA from bodies and heads of both sexes. The position schemes of uORFs and partial CDSs were denoted at the bottom. (C) Dual-luciferase assays conducted on WT and uORF-mutated 5’ UTRs of *Clk* and *cyc* genes in *Drosophila* S2 cells, with reporter constructs depicted top left. For the uORF mutants, the start codons of *Clk* uORF2 and uORF3, as well as *cyc* uORF2, uORF3, and uORF4, were changed from ATGs to ATTs. Renilla luciferase activity was normalized against firefly luciferase and the relative activity in WT was normalized to 1. Six replicates were conducted for each experiment. Data represent mean ± SEM (Two-tailed Student’s *t*-test; ***, *P* < 0.001). (D) Diagrams depicting WT and two uORF knockout strains (*Clk*-uORF-KO and *cyc*-uORF-KO), created using CRISPR-Cas9. For the uORF mutants, the start codons of *Clk* uORF2 and uORF3, as well as *cyc* uORF2, uORF3, and uORF4, were simultaneously changed from ATGs to ATTs. Below each sequence, uORFs (red boxes) and partial CDSs (blue boxes) are shown.

To further explore the regulatory impact of these uORFs on CDS translation, we performed a dual-luciferase reporter assay *in vitro*. We engineered dual-luciferase constructs with either wild-type (WT) or mutated 5’UTRs of *Clk* and *cyc* mRNAs, where the start codons for *Clk* uORF2 and uORF3, as well as *cyc* uORF2, uORF3, and uORF4, were mutated from ATGs to ATTs (Fig. 2C). In *Drosophila* S2 cells, we observed that the luminescence intensity, indicative of CDS translational activity, was more than 70% higher in constructs with mutated 5’UTRs compared to WT controls (WT vs. mutant: 1 vs. 1.79 and 1 vs. 1.83 for *Clk* and *cyc* 5’UTR, respectively; *P* < 0.001 for 5’UTRs of both *Clk* and *cyc*; Fig. 2C). This significant increase confirms the repressive function of these uORFs on the translation of downstream CDSs.

To investigate the function of *Clk* and *cyc* uORFs *in vivo*, we knocked out *Clk* uORF2 and uORF3 simultaneously by mutating their start codon ATGs to ATTs, and established a homozygous knock-out line (*Clk-*uORF-KO) (Fig. 2D). Similarly, we mutated the ATG start codons of uORF2-4 to ATTs in *cyc* and generated a homozygous knock-out line (*cyc-*uORF-KO) (Fig. 2D). We carried out ribosome fractionation followed by quantitative PCR (qPCR) to compare the mRNA abundance in the polysome fraction to that in the monosome fraction (P-to-M ratio) for *Clk* and *cyc*, respectively (Fig. S3). A larger P-to-M ratio means more mRNAs are enriched in the polysome fractions and bound by more ribosomes, thus indicative of higher translation efficiency ^44–46^. We conducted experiments on male heads from *Clk*-uORF-KO, *cyc*-uORF-KO, and WT flies at various Zeitgeber times (ZT), starting from ZT0 (the onset of lights) to ZT20 at 4-hour intervals. P-to-M ratios of *Clk* mRNA in *Clk*-uORF-KO flies significantly exceeded WT at ZT0, ZT4, and ZT8, with no notable changes at ZT12 and ZT16, and a decrease at ZT20 (Fig. 3A). Conversely, P-to-M ratios for *cyc* mRNA in *cyc*-uORF-KO flies were slightly higher at all but one ZT time point (ZT12) (Fig. 3B). Notably, the elevation in *cyc* mRNA was not statistically significant except at ZT16. Overall, these patterns indicate *Clk* uORFs exert a substantial impact on translation, especially during the light phase, while *cyc* uORFs show modest effects. Consequently, our research has pivoted to focus on *Clk* uORFs.

**Fig. 3.**
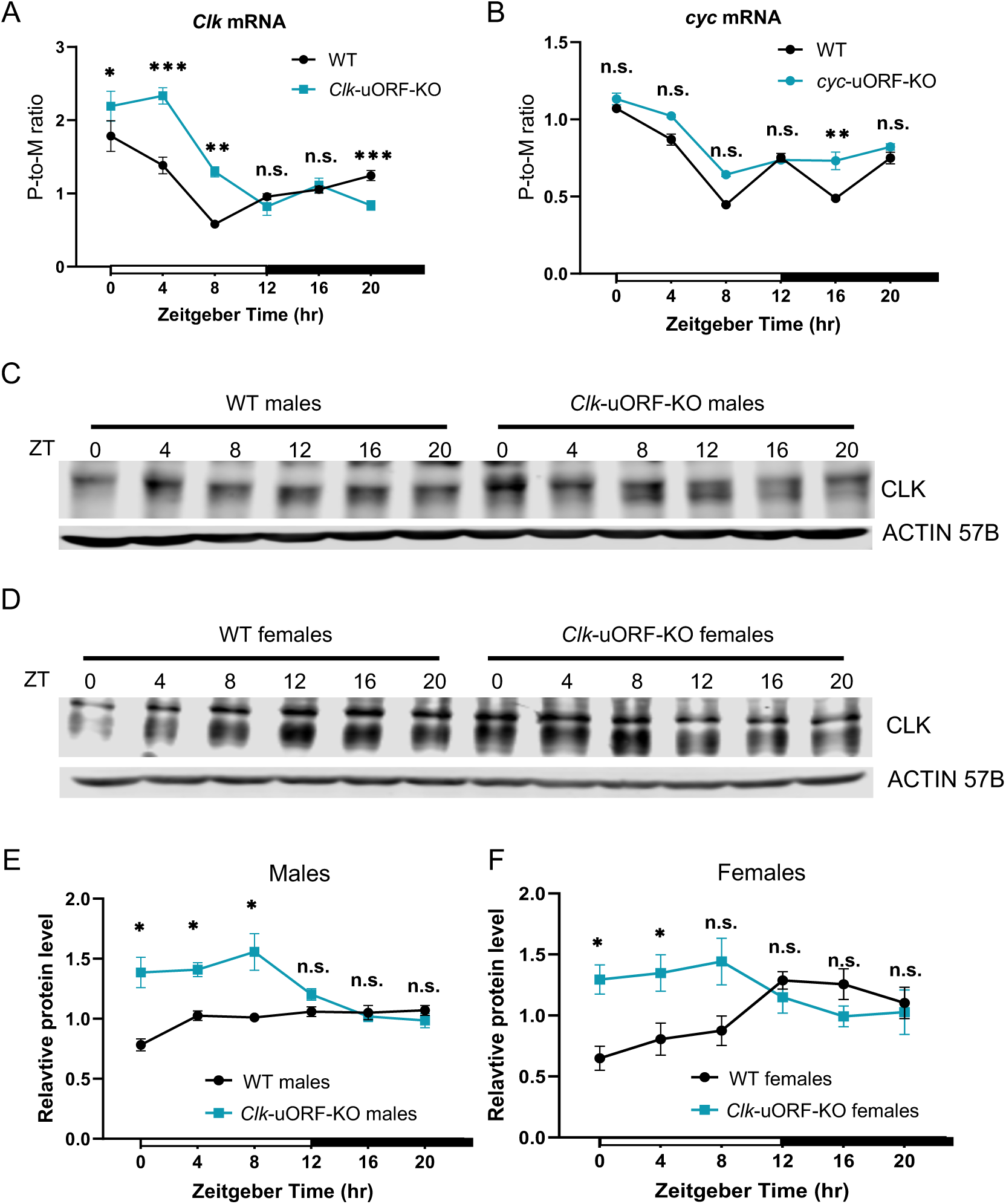
*Clk* uORFs suppress CLK protein translation in fly heads *in vivo*. (A) P-to-M ratios of *Clk* mRNA from whole heads of male *Clk*-uORF-KO compared to that in male WT flies, sampled at indicated Zeitgeber times at 4-hour intervals. Each time point includes six biological replicates. Data are expressed as mean ± SEM (Wilcoxon signed-rank test. *, *P* < 0.05; **, *P* < 0.01; ***, *P* < 0.001; n.s., *P* > 0.05). (B) P-to-M ratios of *cyc* mRNA from whole heads of male *cyc*-uORF-KO compared to that in male WT flies, sampled at indicated Zeitgeber times at 4-hour intervals. Each time point includes six biological replicates. Data are expressed as mean ± SEM (Wilcoxon signed-rank test. **, *P* < 0.01; n.s., *P* > 0.05). (C) Representative Western blots of CLK protein from whole-head protein extracts of male flies, sampled at indicated Zeitgeber times at 4-hour intervals. ACTIN 57B served as the loading control. (D) Representative Western blots of CLK protein from whole-head protein extracts of female flies, sampled at indicated Zeitgeber times at 4-hour intervals. ACTIN 57B served as the loading control. (E) Quantification of CLK abundance relative to ACTIN 57B from (C), with the average intensity for WT normalized to 1 at each Zeitgeber Time (ZT) point. The analysis included four replicates. Data are presented as mean ± SEM (Two-tailed Mann-Whitney *U* test. *, *P* < 0.05; n.s., *P* > 0.05). (F) Quantification of CLK abundance relative to ACTIN 57B from (D), with the average intensity for WT normalized to 1 at each Zeitgeber Time (ZT) point. The analysis included four replicates. Data are presented as mean ± SEM (Two-tailed Mann-Whitney *U* test. *, *P* < 0.05; **, *P* < 0.01; n.s., *P* > 0.05).

Next, we measured CLK protein abundances for male and female heads at different time points using immunoblotting (Fig. 3C and 3D). We found that CLK protein levels were significantly higher in the heads of male *Clk*-uORF-KO flies compared to WT during the morning hours of ZT0, ZT4, and ZT8, and marginally lower, though not significantly, at ZT16 and ZT20 (Fig. 3E). Similar patterns were observed for female heads (Fig. 3F). These protein abundance trends corroborate the enhanced translational activity in *Clk*-uORF-KO mutants compared to WT during the light phase. Overall, the data demonstrate that *Clk* uORFs downregulate CLK protein translation during the daytime, with pronounced effects observed in the early hours.

### *Clk* uORFs modulate the pace of the circadian clock

To quantitatively assess the effects of *Clk* uORF KO on circadian rhythms, we utilized a mathematical model previously established for simulating the *Drosophila* molecular clock ^47^. This model employs nonlinear ordinary differential equations to model the regulatory weights among mRNA and protein molecules within the circadian network, making it applicable for predicting the increase in *Clk* translation when uORFs are disrupted. By incrementing the regulatory weight from *Clk* mRNA to CLK protein from the baseline to higher values, we incorporated the anticipated enhancement in *Clk* translation due to the uORF KO into our model (Fig. 4A).

**Fig. 4.**
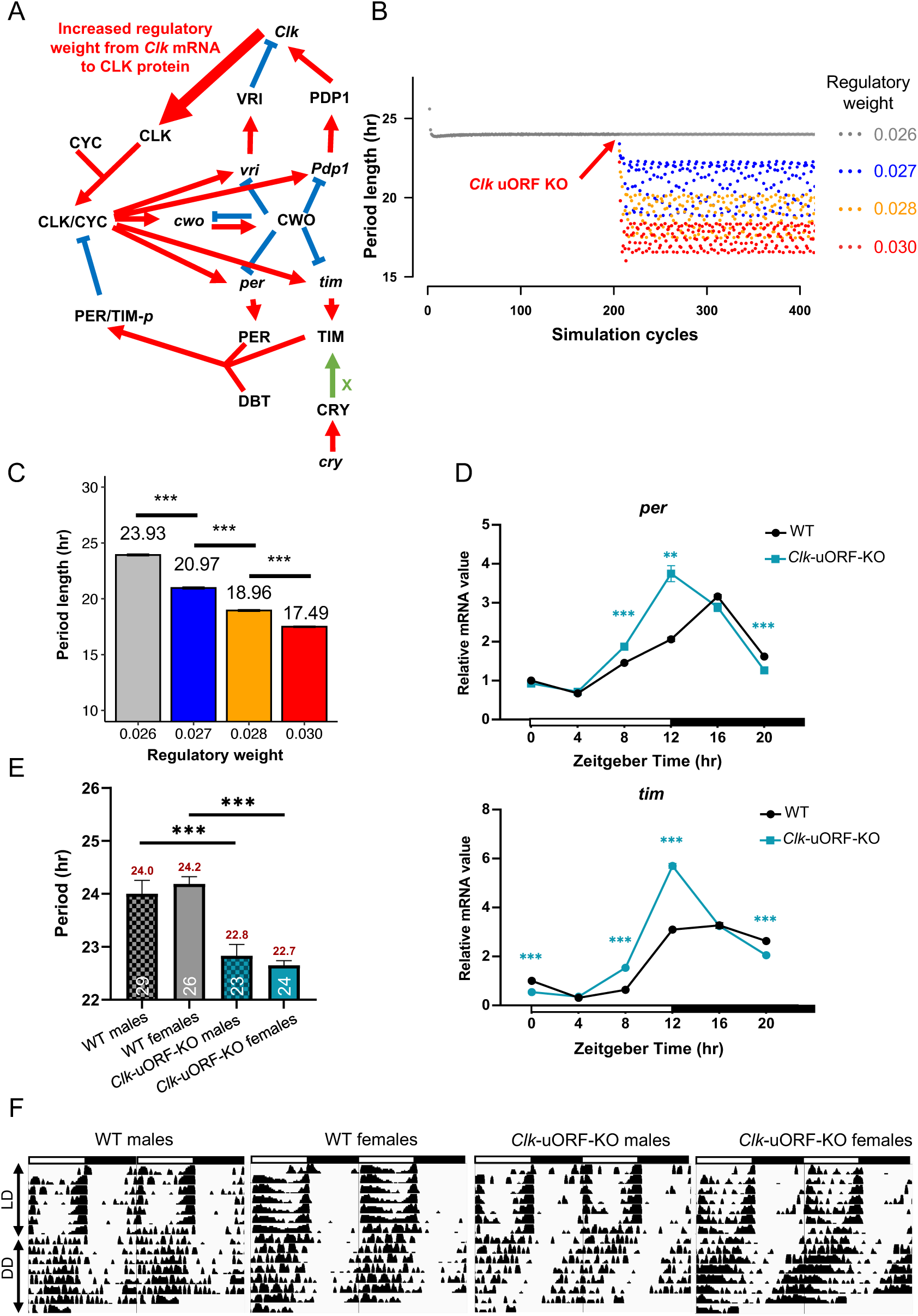
Computational and experimental analysis of *Clk* uORFs on circadian rhythms. (A) Diagram of the *Drosophila* circadian clock molecular network, adapted from a previous study ^47^. mRNA molecules are indicated in italicized lowercase, while protein molecules are in uppercase. Red arrows represent activation, while blue lines indicate suppression. The bold red arrow from *Clk* mRNA to CLK protein signifies the enhanced translation activity due to *Clk* uORF knockout. The green arrow with an “X” indicates that CRY protein promotes the degradation of TIM. (B) The distribution of period length (hr) under different *Clk* mRNA to CLK protein regulatory weights across approximately 400 simulation cycles. The original regulatory weight setting of 0.026 in the original model ^47^ acts as the baseline, emulating the presence of WT uORF. After 200 cycles the regulatory weight is increased from 0.0026 to several arbitrary higher levels (0.027, 0.028, and 0.029) which is denoted by the red arrow, and continued for another 200 simulation cycles. (C) The statistics of circadian period length (hr) for simulations in (B). Data are measured in 200 simulation cycles and are reported as the mean ± SEM (two-tailed Student’s *t* test. ***, *P* < 0.001). (D) qRT-PCR measured mRNA levels of *per* and t*im* from total RNA extracts of female heads, sampled at specified Zeitgeber time points. WT intensity at ZT0 is normalized to 1. Each time point includes four biological replicates. Data are reported as the mean ± SEM (two-tailed Student’s *t* test. **, *P* < 0.01; ***, *P* < 0.001). (E) Period of locomotor rhythms of *Clk*-uORF-KO flies and WT flies under constant darkness conditions (DD). The period lengths (hr) and the number of flies tested (*n*) are displayed above and below each bar, respectively. (F) Representative double-plotted actograms of the locomotor activity of the indicated flies, monitored under LD followed by DD. White and black bars represent light and dark phases, respectively.

The simulation indicated an immediate increase in CLK protein levels correlating with a higher regulatory weight from *Clk* mRNA to protein, which mimics the effect of *Clk* uORF KO (Fig. S4A). This, in turn, led to elevated levels of *per* and *tim* mRNA and protein (Fig. S4A and S4B). In the simulations, we analyzed the intervals between peaks of *tim* mRNA, using these as a measure of circadian period length. The mean ± SEM of period length is 23.926 ± 0.003 hours, 20.967 ± 0.078 hours, 18.956 ± 0.061 hours, and 17.494 ± 0.043 hours for regulatory weights from *Clk* mRNA to CLK protein set at 0.026 (baseline), 0.027, 0.028, and 0.030, respectively (Fig. 4B and 4C; SI Appendix, Table S2). These findings demonstrate a shortening of the circadian period as the regulatory weight increases, suggesting that enhanced *Clk* translation—resulting from the disruption of *Clk* uORFs—compresses the circadian period.

To corroborate the simulation outcomes, we measured the mRNA level of *per* and *tim*, observing significant elevation in *Clk-*uORF-KO flies with the peak level occurring at an earlier time in female heads (Fig. 4D). As our simulations predicted, disrupting *Clk* uORFs led to a reduced period length by over one hour for the locomotor rhythm of *Clk*-uORF-KO flies. Specifically, the period’s mean ± SEM for WT males is 24.0 ± 0.25, WT females 24.2 ± 0.14, *Clk*-uORF-KO males 22.8 ± 0.21, and *Clk*-uORF-KO females 22.7 ± 0.09 (Fig. 4E and 4F; SI Appendix, Table S3). These empirical results, together with our simulation data, strongly indicate that *Clk* uORFs modulate CLK translation, thereby regulating the timing of the molecular clock through the expression of downstream genes like *per* and *tim*.

### *Clk* uORFs influence the temporal transcriptomic landscape

CLK modulates the cyclical expression of a significant portion of *Drosophila* transcriptome ^48^. To explore the impact of *Clk* uORFs on the global gene expression rhythms, we harvested heads from WT and *Clk*-uORF-KO flies at four-hour intervals across the day, using three biological replicates, and performed RNA sequencing for transcriptome analysis. Using JTK-CYCLE ^49^ to detect genes with oscillating mRNA levels (Table S4), we found 3,370 and 3,859 genes exhibited rhythmic expression in WT and *Clk*-uORF-KO female heads, respectively, with 2,235 genes in common (Fig. 5A and 5B). In male heads, 3,570 genes showed rhythmic expression in WT, whereas 3,890 were rhythmic in *Clk*-uORF-KO, with 2,124 genes in common (Fig. 5C; SI Appendix, Fig S5). Overall, knocking out *Clk* uORFs disrupted the rhythmic expression of about 34-41% of genes in WT, while approximately 1,700 genes displayed *de novo* cyclic expression patterns in *Clk*-uORF-KO heads. Gene ontology (GO) analysis demonstrated that genes which lost rhythms in *Clk-*uORF-KO flies were mainly involved in rRNA processing, ribosome biogenesis, translation and metabolism (SI Appendix, Fig. S6A and S6C), while genes that gained rhythms when *Clk* uORFs were knocked out were enriched in pathways such as translation, mitochondrial processes and development (SI Appendix, Fig. S6B and S6D).

**Figure 5.**
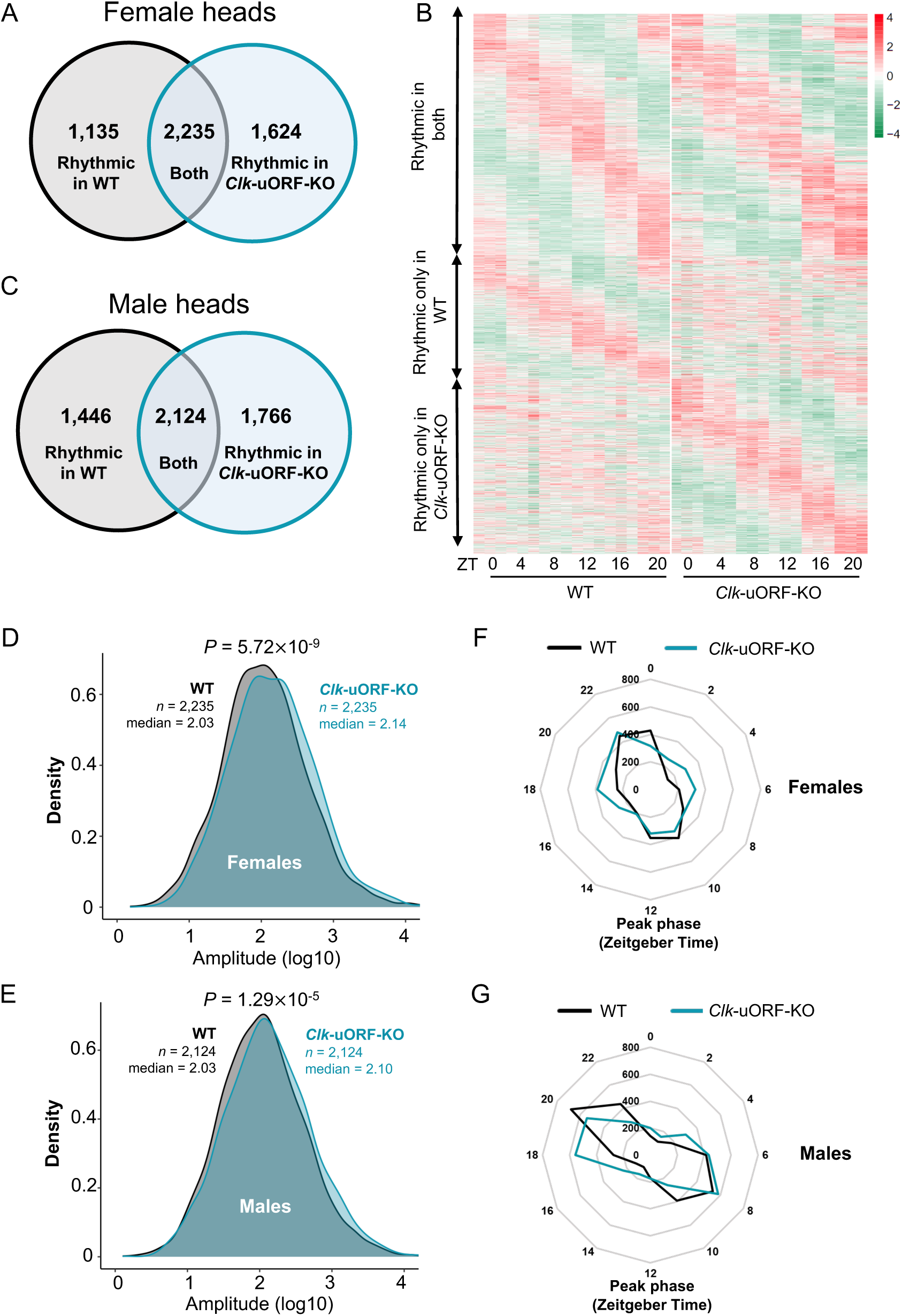
*Clk* uORFs modulate the temporal pattern of transcriptome expression landscape in fly heads. (A) The number of rhythmically expressed genes identified from whole-head RNA-seq data of female WT and *Clk*-uORF-KO flies. (B) Hierarchical clustering of genes expressed rhythmically in whole heads of female flies across various Zeitgeber time points, with gene expression levels normalized to Z-scores. The color scale reflects expression intensity, ranging from low (green) to high (red) Z-scores. (C) The number of rhythmically expressed genes identified from whole-head RNA-seq data of male WT and *Clk*-uORF-KO flies. (D) The distribution of expression amplitudes for genes rhythmically expressed in the heads of female WT and *Clk*-uORF-KO flies (*n* = 2,235; Wilcoxon rank sum test; *P* = 5.72×10^-9^). (E) The distribution of expression amplitudes for genes rhythmically expressed in the heads of male WT and *Clk*-uORF-KO flies (*n* = 2,124; Wilcoxon rank sum test; *P* = 1.29×10^-5^). (F-G) Phase distribution of rhythmically expressed genes in whole heads of female WT and *Clk*-uORF-KO flies (F) and male WT and *Clk*-uORF-KO flies (G). The *y*-axis of the radar chart indicates the number of rhythmic genes.

Remarkably, genes related to circadian rhythms were significantly overrepresented among rhythmically expressed genes in *Clk*-uORF-KO female heads (86 out of 150) compared to the rest of the expressed genes (3,773 out of 9,051, Fisher’s exact test, *P* = 0.02). This enrichment was more significant in the heads of *Clk*-uORF-KO females than in WT females, WT males, or *Clk*-uORF-KO males. Knocking out *Clk* uORFs led to an increase in oscillation amplitude which was more significant in females than in males (Wilcoxon rank sum test; *P* = 5.72×10^-9^ in females vs. *P* = 1.29×10^-5^ in males; Fig. 5D and 5E). Furthermore, in females, knocking out *Clk* uORFs led to desynchronized oscillations with peak expression time distributed throughout the day (Fig. 5F and 5G). In summary, disrupting *Clk* uORFs markedly influences gene expression rhythms in fly heads, exerting more pronounced effects in females than males. These alterations in gene expression could result in diverse impacts on physiological functions, especially in females, aspects of which are investigated in the following sections.

### *Clk* uORFs promote morning wakefulness by tuning up the dopaminergic tone

Building on evidence that CLK promotes sleep ^50^, we first examined the impact of knocking out *Clk* uORFs on sleep. We found that knocking out *Clk* uORFs significantly lengthened sleep duration, with more prominent alteration in females (9.5% increase; Fig. 6A; SI Appendix, Fig. S7A) compared to males (4.0% increase; SI Appendix, Fig. S7B and S7C). A closer examination showed that in female *Clk*-uORF-KO flies, the duration of sleep was notably prolonged by 38.65% during the initial 8 hours of the light phase, indicating that *Clk* uORFs facilitate morning wakefulness (Fig. 6B). The increase in sleep duration stems from a longer average sleep bout, implicating enhanced sleep continuity upon *Clk* uORF deletion (Fig. 6C). Additionally, waking activity levels were higher in female *Clk*-uORF-KO flies, indicating that the prolonged sleep is not due to impaired motor function (Fig. 6C). While in males, the alterations in sleep were not as prominent as that observed in females (SI Appendix, Fig. S7D).

**Fig. 6.**
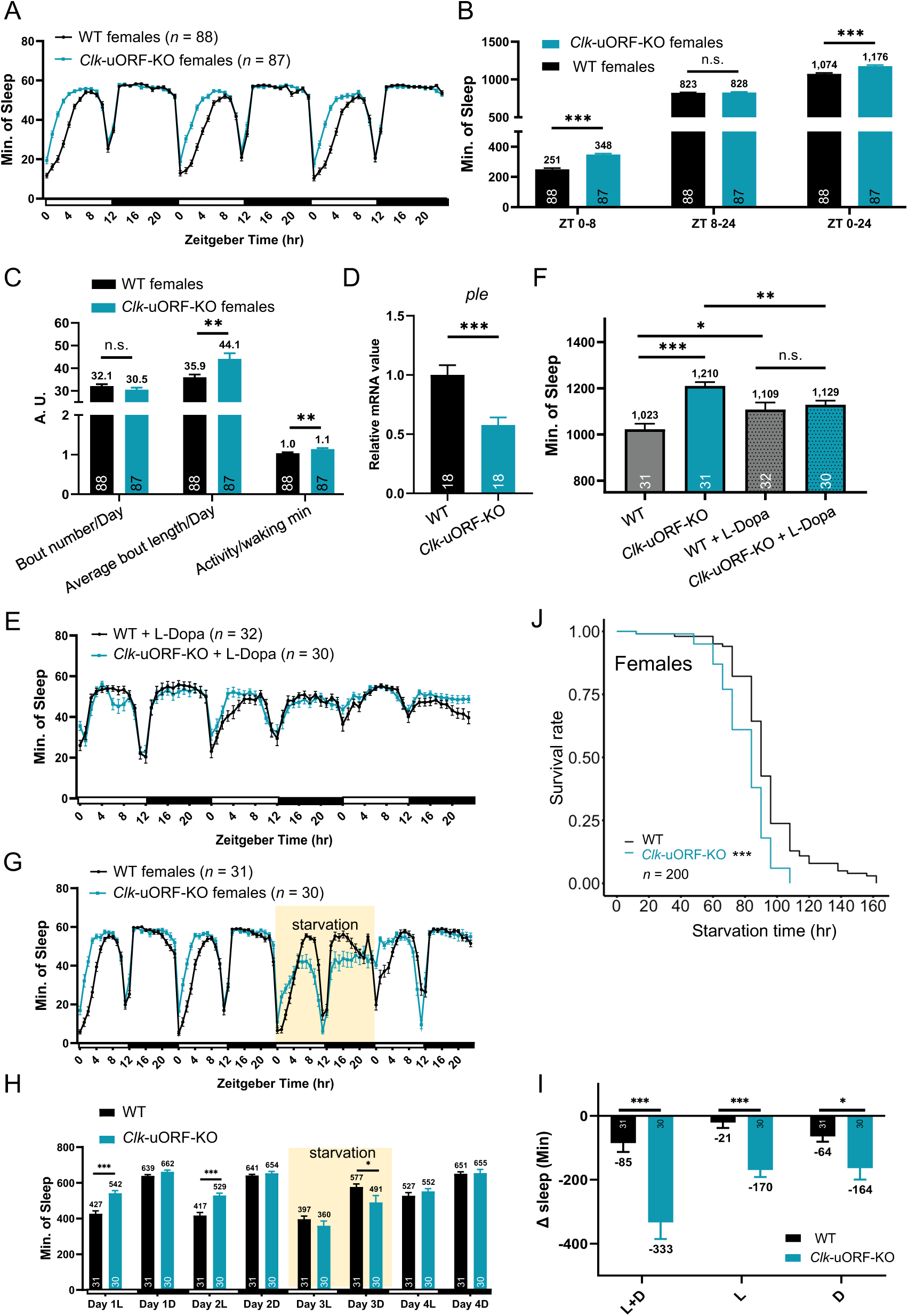
*Clk* uORFs regulate sleep/wakefulness. (A) Sleep profile of female *Clk-*uORF-KO and WT flies under LD conditions. (B) Sleep duration during ZT0-8, ZT8-24, and ZT0-24 of female *Clk-*uORF-KO and WT flies under LD condition. (C) Daily sleep bout number, average bout length, and average waking activity of female *Clk-*uORF-KO and WT flies under LD condition. (D) The relative mRNA abundance of *ple* determined by RT-qPCR using whole-head total RNA extracts collected from female flies under LD conditions. The average value of WT is set to 1. (E) Sleep profile of female *Clk-*uORF-KO and WT flies treated with L-Dopa under LD condition. (F) Daily sleep duration of female *Clk-*uORF-KO *vs.* WT flies treated with L-Dopa. (G) Sleep profile of female uORF-KO and WT flies under food ad libitum and starvation (the 3^rd^ day) condition. The yellow shade indicates period of starvation. (H) Daily sleep duration of female *Clk-*uORF-KO and WT flies under baseline and starvation conditions. (I) Histogram shows starvation-induced sleep change (Δ sleep) compared with normal conditions during daytime (L), nighttime (D), and the entire day (LD). (J) Survival curves of female flies under starvation conditions (*n* = 200; log-rank test; ***, *P* < 0.001). The numbers of flies tested are shown in the brackets of legend or the bottom of histograms. Data are presented as the mean ± SEM. Asterisks indicate statistical significance (two-tailed Student’s *t* test. *, *P* < 0.05; **, *P* < 0.01; ***, *P* < 0.001; n.s., *P* > 0.05). ZT, Zeitgeber Time.

Given that CLK suppresses expression of the gene encoding tyrosine hydroxylase (*ple*), which in turn diminishes dopamine signaling and activity levels ^51^, we investigated whether dopamine signaling mediates the influences of *Clk* uORFs on sleep. Our qPCR assays revealed a notable decrease in *ple* mRNA in female *Clk*-uORF-KO heads compared to WT (Fig. 6D), aligning with the hypothesis that elevated CLK protein levels—resulting from *Clk* uORF disruption—downregulate *ple* expression. Next, we treated female flies with dopamine precursor L-DOPA, which facilitates dopamine synthesis ^52^. This treatment reversed the prolonged sleep duration in female *Clk-*uORF-KO flies (Fig. 6E and 6F). Thus, *Clk* uORFs appear to promote morning alertness by modulating dopaminergic signaling.

### *Clk* uORFs limit starvation-induced wakefulness and are beneficial for survival

Starvation typically reduces sleep through a mechanism dependent on CLK, a response thought to enhance survival chances ^53^. To determine if *Clk* uORFs modulate the effects of starvation on sleep/wakefulness, we analyzed sleep patterns in female *Clk*-uORF-KO flies during starvation. We observed an exaggerated sleep loss in *Clk-*uORF-KO flies, suggesting that *Clk* uORFs may function to restrict starvation-induced sleep loss (Fig. 6G-I). Moreover, the increased wakefulness in *Clk*-uORF-KO flies correlated with significantly faster mortality rates under starvation compared to WT flies for both sexes [*P* = 8.8×10^-13^ for females (Fig. 6J) and *P* = 2.4×10^-9^ for males (SI Appendix, Fig. S8), log-rank test]. These results indicate that *Clk* uORFs function to restrict sleep loss during starvation, which potentially affects survival.

### *Clk* uORFs modulate sleep/wakefulness in adaptation to seasonal photoperiodic changes by suppressing CLK protein translation in a photoperiod-dependent manner

Given the known significance of the circadian clock in adapting to seasonal environmental variations ^54^, we hypothesized that *Clk* uORFs might similarly regulate translation on a circannual basis. To test this hypothesis, we subjected female *Clk-*uORF-KO and WT flies to conditions that mimic seasonal changes. Because the most important seasonal signal for the circadian clock is the shortening/lengthening of photoperiod ^54^, we monitored locomotor rhythm and sleep under photoperiods of different lengths, ranging from 4 hours of light and 20 hours of dark (4L20D) to 20 hours of light and 4 hours of dark (20L4D). Our observations indicate that *Clk* uORFs did not influence the timing of locomotor activity rhythm across these varied photoperiods, suggesting that *Clk* uORFs are not essential for adapting locomotor rhythms to changes in seasonal day length (SI Appendix, Fig. S9).

Nevertheless, analysis of sleep duration under varying photoperiods revealed that female *Clk*-uORF-KO flies exhibited a greater disparity from WT as the photoperiod decreased (Fig. 7A and 7B). Notably, sleep duration showed a significant positive correlation with photoperiod length in WT (Pearson’s *r* = 0.323, *P* = 6.5 × 10^-8^), while this correlation was much weaker in *Clk*-uORF-KO (Pearson’s *r* = 0.172, *P* = 0.006) (Fig. 7C). As the consequence, the difference in sleep duration between *Clk*-uORF-KO and WT became larger as the photoperiod shortened (Fig. 7B). Under 4L20D, the shortest photoperiod tested here, WT displayed on average 918 ± 26 minutes of sleep, while that of *Clk*-uORF-KO females was 1,124 ± 18 minutes, with an increase of 206 minutes on average (22.44%). In contrast, under 20L4D, the longest photoperiod tested, WT females slept for 1,082 ± 21 minutes compared to 1,195 ± 18 minutes for *Clk*-uORF-KO females, yielding a smaller difference of 113 minutes on average (10.44%). These findings indicate that *Clk*-uORF-KO flies showed a diminished response to changes in photoperiod length, implying that *Clk* uORF-mediated translational regulation plays a role in synchronizing sleep patterns with seasonal daylight variations.

**Fig. 7.**
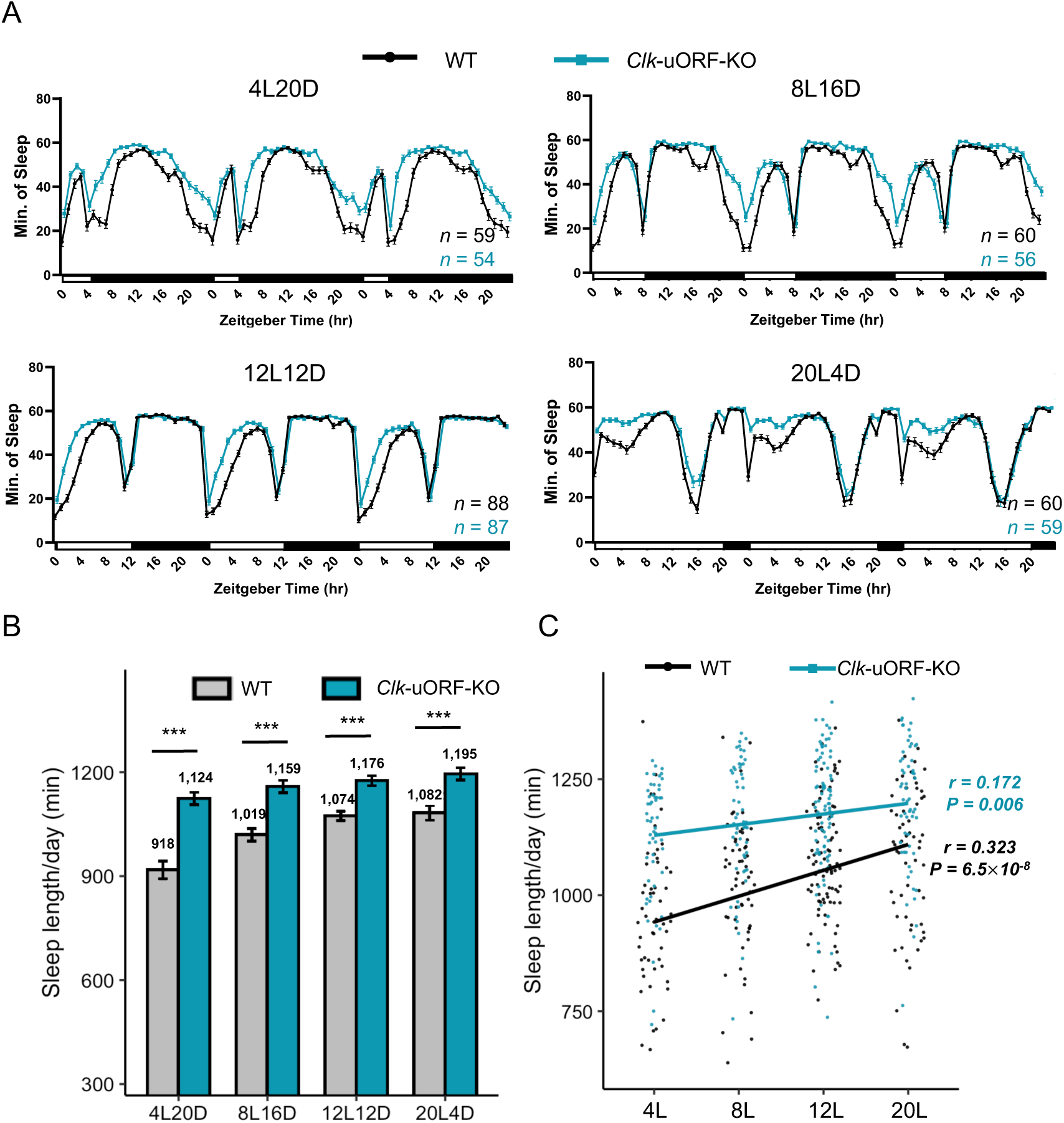
*Clk* uORFs promote wakefulness in adaptation to seasonal photoperiod changes. (A) Sleep profile of female *Clk-*uORF-KO and WT flies under 4 h L : 20 h D (4L20D) (A), 8 h L : 16 h D (8L16D) (B), 12 h L : 12 h D (12L12D) (C), and 20 h L : 4 h D (20L4D) (D) condition. The numbers of tested flies (*n*) are shown in the bottom-right. (B) Daily sleep duration (min) of female *Clk-*uORF-KO and WT flies under different photoperiods. Data are presented as the mean ± SEM. Asterisks and dollar signs indicate statistical significance (two-tailed Student’s *t* test. ***, *P* < 0.001). (C) The correlation between daily sleep duration (min) and photoperiod length in female *Clk-*uORF-KO and WT flies (Pearson’s correlation).

Given these observed behavioral changes, we posited that *Clk* uORFs participate in translational regulation in a photoperiod-dependent manner. To test this, we carried out ribosome fractionation followed by qPCR under 4L20D and 12L12D light-dark cycles. In WT heads, we observed a consistent decrease in P-to-M ratios throughout the day under 12L12D vs. 4L20D (Fig. 8A). However, such a pattern was not observed in *Clk-*uORF-KO (Fig. 8B). The area under the curve (AUC) of these ratios, indicative of overall daily CLK protein synthesis, was significantly reduced by long photoperiods in WT fly heads (∼30% reduction), while the reduction was much less (∼8% reduction) in *Clk*-uORF-KO (Fig. 8C and 8D). This photoperiod dependency was corroborated at the protein level. In WT heads, CLK protein levels dipped significantly during ZT4-12 under 4L20D compared to 12L12D (Fig. 8E and 8F). In contrast, CLK protein level was only significantly decreased at ZT0 in *Clk-*uORF-KO under 4L20D relative to 12L12D (Fig. 8E and 8F). Moreover, the AUC analysis demonstrated that short photoperiod significantly reduced total daily CLK level in WT but not in *Clk-*uORF-KO heads (SI Appendix, Fig. S10). Collectively, these findings suggest that *Clk* uORFs are involved in photoperiod-dependent translation suppression, which likely aids in adjusting wakefulness to seasonal variations.

**Fig. 8.**
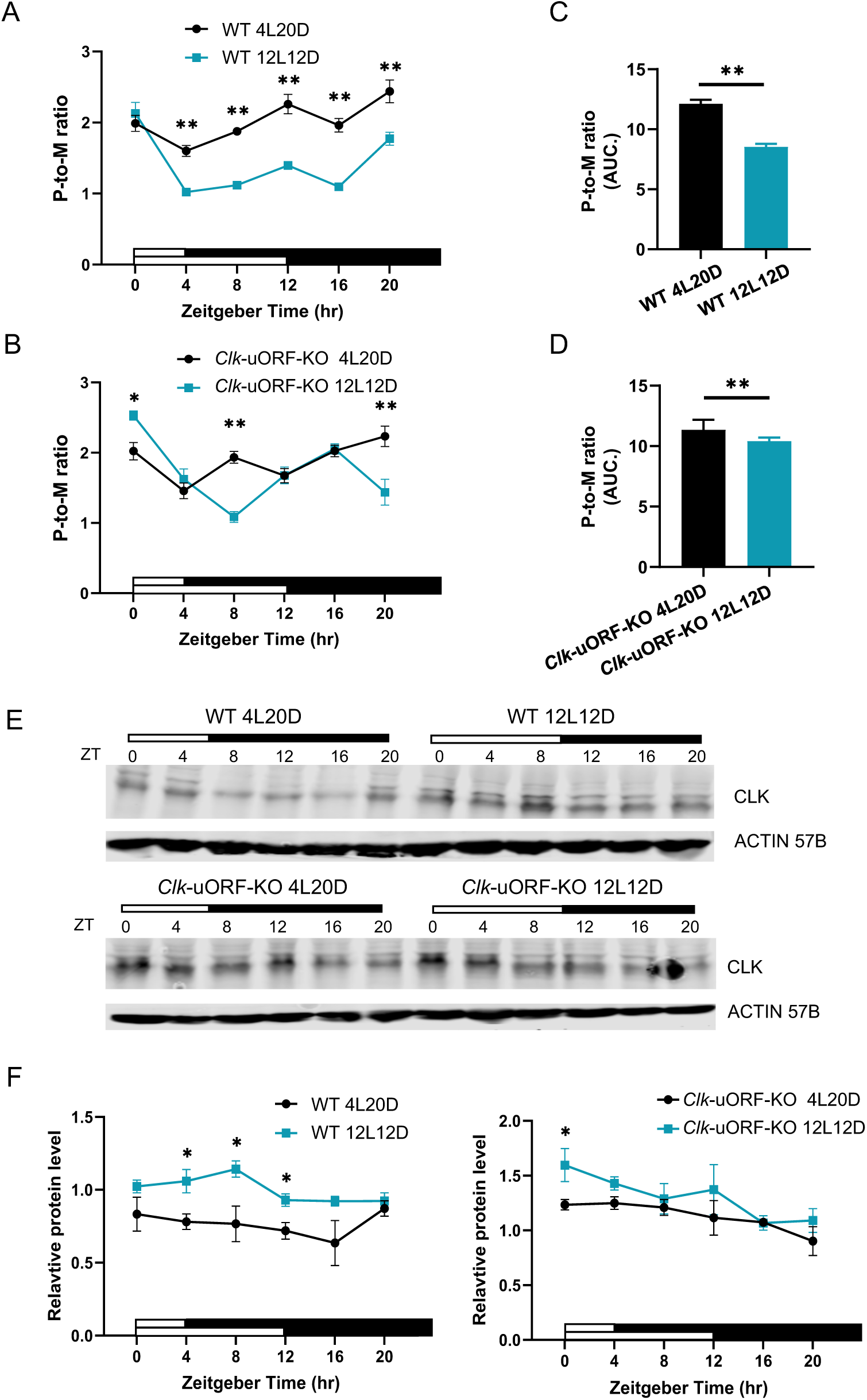
*Clk* uORFs suppress CLK protein translation in a photoperiod-dependent manner. (A-B) P-to-M ratios of *Clk* mRNA from whole-head extracts collected from female WT (A) and *Clk-*uORF-KO (B) flies at indicated time points under 4L20D and 12L12D conditions. Each time point includes six biological replicates and data are presented as the mean ± SEM. (C-D) AUC analysis of daily P-to-M ratio in (A) and (B) of female WT (C) and *Clk-*uORF-KO (D) flies under indicated photoperiod. (E) Representative Western blots for CLK protein from whole-head total protein extracts collected from female WT (upper) and *Clk-*uORF-KO (lower) flies at indicated time points under 4L20D (left) and 12L12D (right). ACTIN 57B is used as a loading control. (F) The relative CLK abundance normalized to ACTIN 57B in (E). The average value of WT under 12L12D is set to 1. Each time point includes four biological replicates and data are presented as the mean ± SEM. Asterisks indicate statistical significance (two-tailed Mann-Whitney *U* test for unpaired comparisons. **, *P* < 0.01; ***, *P* < 0.001). ZT, Zeitgeber Time.

### Knocking out *Clk* uORFs impairs fecundity

Recognizing the roles of *Clk* uORFs in facilitating the adaptation to environmental changes, we further investigated whether they enhance evolutionary fitness by assessing fecundity. Remarkably, mated *Clk*-uORF-KO females laid significantly fewer eggs than their WT counterparts during an eight-day monitoring duration, with the total egg production of mutants markedly less (*P* < 0.01) (SI Appendix, Fig. S11A). This pattern persisted in *Clk*-uORF-KO virgin females, which also laid fewer eggs than WT in both segmented (2-days) and pooled (10-days) measurements (SI Appendix, Fig. S11B). Accordingly, female *Clk-*uORF-KO flies produced significantly fewer offspring than WT counterparts at 25°C (*P* = 1.5×10^-5^ Wilcoxon rank sum test) (SI Appendix, Fig. S11C). Even at 29°C, when both mutants and WT flies exhibited reduced offspring production compared to 25°C, the offspring counts for *Clk-*uORF-KO females were still significantly fewer than that for WT (*P* = 0.00029, Wilcoxon rank sum test) (SI Appendix, Fig. S11C). Collectively, these results indicate that *Clk* uORFs profoundly affect various aspects of physiological and behavioral processes, with implications for fitness beyond their established roles in circadian rhythm and sleep regulation.

## Discussion

The evolutionary conservation of the circadian mechanisms across species underscores their fundamental significance. Our understanding of circadian rhythms has grown substantially over the years, providing critical insights into a range of physiological processes. Earlier investigations, including our own, have noted a very low correlation between the temporal variation pattern of the transcriptome and that of the proteome, indicating that regulations at the translational and post-translational levels play dominant roles in determining the daily variations of protein abundance ^55^. Within this complex regulatory network, our study here has highlighted a pivotal role for translational regulation, specifically mediated by uORFs, in fine-tuning these rhythms. Beyond circadian rhythm and sleep, *Clk-*uORF-KO flies show diminished fecundity and reduced starvation resilience, suggesting that *Clk* uORFs might act as a master regulator, shaping the rhythmic expression of a vast array of genes and influencing multifaceted physiological outcomes.

In this study, we observed over one-hour shortening of the circadian period when *Clk* uORFs were knocked out, which means these uORFs play a crucial role in the core circadian timing mechanism. Consistent with this idea, we found that *Clk* uORFs mediate an instructive signal to the molecular clock by suppressing CLK protein translation during the daytime. Besides setting the pace of the clock, *Clk* uORFs also modulate cyclic gene expression globally. The oscillation of over 1/3 of the rhythmic genes requires *Clk* uORFs, as these oscillations were dampened in *Clk-*uORF-KO flies. This reveals the delicacy of the circadian system: an increase of CLK translation and, thus, CLK protein level caused by knocking out *Clk* uORFs can exert detrimental effects on the rhythmicity of the transcriptome. It is surprising that genes cycle with a more robust amplitude in *Clk-*uORF-KO flies relative to WT. Among these genes, about half did not display oscillatory expression patterns in WT. In other words, *Clk* uORFs suppress the mRNA rhythms of these genes. We reason that knocking out *Clk* uORFs increases CLK protein level in a time-dependent manner, which results in enhanced transcription of CLK target genes at a certain time of the day, and thus, these genes may acquire a *de novo* rhythmic expression pattern. Considering that *Clk* uORFs suppress CLK translation more under longer photoperiods and less under shorter photoperiods, we speculate this photoperiod-dependent variation in the translational activities of *Clk* uORFs can shape the rhythmic transcription landscape in adaptation to seasonal photoperiod changes.

*Clk* uORFs promote morning wakefulness, consistent with the time-dependent modulation of these uORFs on CLK protein translation and abundance. This regulation is not only time of the day-dependent, but also time of the year-dependent. WT flies display increased wakefulness and decreased sleep in adaptation to a shortening of photoperiod (occurring in fall and winter seasons), while *Clk* uORFs participate in modulating this process by suppressing CLK translation in a photoperiod-dependent manner. Intriguingly, human individuals with seasonal affective disorder (SAD, commonly known as winter depression) report a winter increase in sleep, ranging from 30 min to 2 h longer in duration compared to controls, while the underlying mechanism is yet unknown ^56^. The long sleep phenotype caused by knocking out *Clk* uORFs is most striking under a short photoperiod, reminiscent of the winter hypersomnolence in SAD. Notably, while we observed a stronger sleep phenotype in female flies, SAD is also more predominant in women ^57^. From a mechanistic perspective, our results strongly suggest that *Clk* uORFs promote wakefulness by increasing tyrosine hydroxylase expression, thus up-regulating the dopaminergic tone. Similarly, the mammalian CLOCK protein has been shown to repress the transcription of *Tyrosine hydroxylase*, while *Clock* mutation leads to decreased sleep and depression-like behaviors in mice due to elevated dopamine signaling ^58,59^. Taken together, our findings here may provide some insights for understanding the pathological mechanism of SAD.

It seems contradictory that CLK translation is down-regulated under 12L12D vs. 4L20D, while CLK protein level is higher under 12L12D relative to 4L20D. We reason that CLK protein abundance is determined by coordinated actions of transcription, translation, and post-translational modifications. A longer photoperiod may promote transcription and/or inhibit degradation of CLK protein, while *Clk* uORFs counteract these influences by suppressing CLK synthesis. In other words, *Clk* uORFs may serve as a buffer against the impacts of transcription and/or PTMs, adjusting CLK protein abundance to maintain an optimal level throughout the year. Of course, further investigations are needed to elucidate how *Clk* uORFs mediate photoperiodic signals.

uORFs have been shown to dynamically modulate the translation of CDSs in stress responses ^30^, and here we also observed a role for *Clk* uORFs in responses to starvation stress. Although under food *ad libitum* condition, *Clk* uORFs promote wakefulness, under starvation they function to reduce wakefulness, implicating the complexity of uORF regulation and their multifaceted roles in varying physiological states. Despite these complicated and sometimes contradictory actions, in the end, the presence of *Clk* uORFs confers adaptive advantages and is beneficial for the survival (KO flies die earlier when starved) and fecundity of the organism.

To summarize, here we observed an enrichment of uORFs in circadian rhythm genes and further identified functional uORFs in *Clk* that repress the translation of CLK protein in both circadian and circannual manners. These regulations slow down the pace of the clock, modulate global rhythms of gene expression, and promote morning wakefulness in adaption to seasonal photoperiod changes. These findings, situated within the broader evolutionary, clinical, and interdisciplinary contexts, underscore the pivotal role of uORFs in shaping biological processes and potential therapeutic avenues.

## Acknowledgements

This work was supported by grants from the Natural Science Foundation of China (31930021 and 32341021 to L.Z., 32070597 to J. L, and 32200956 to K.S.), the Ministry of Science and Technology of China (STI 2030-Major Projects 2021ZD0203200-02 to L.Z, and 2022YFE0132000 to J.L.), the Department of Science and Technology of Hubei Province (2022CFA049 to L.Z.) and Natural Science Foundation of Beijing (grant 5212006 to J.L.). We thank the National Center for Protein Sciences at Peking University for technical assistance. Some of the analyses were performed on the High-Performance Computing Platform of the Center for Life Sciences. We thank Dr. Joanna Chiu for CLK antibody used in this study.

## Data and code availability

- The raw fastq files for the RNA-seq libraries were deposited to NCBI Sequence Read Archive (SRA) with BioProject accession PRJNA1026336 and are publicly available as of the date of publication.
- Original Western blot images have been deposited to Mendeley and are publicly available on the date of publication.
- Any additional information required to reanalyze the data reported in this paper is available from the lead contact upon request.

## Materials and Methods

### Fly strains and cell line

All flies were reared on standard cornmeal-yeast-sucrose medium and kept in light/dark (LD) cycles at 25°C except for specific experimental design denoted in the text. Food was accessible *ad libitum*. *{nos-Cas9}attP40* and *{nos-Cas9}attP2* fly stocks used for embryo injection were from the TsingHua Fly Center. The tool fly stocks (*y sc v* and *Dr, e/TM3, Sb*) used for mutant screening were from the TsingHua Fly Center*. Clk-*uORF-KO and *cyc*-uORF-KO mutants were generated and backcrossed to the Canton-S strain for nine generations. Homozygous mutants were used throughout this study and the Canton-S strain served as the WT control flies. *Drosophila* S2 cells used in this study were purchased from Life Technologies Corp (http://www.lifetechnologies.com).

### The conservation assessment of uORFs

We retrieved the whole genome sequence alignment (maf) of *D. melanogaster* (dm6) with 22 other *Drosophila* species from the UCSC genome browser (genome.ucsc.edu). Leveraging the genome annotation of *D. melanogaster* (FlyBase r6.04, https://flybase.org/), the 5’UTR sequence of each annotated transcript of *D. melanogaster* and its corresponding sequences in other 22 *Drosophila* were extracted from the maf. We scanned all the AUG triplets within the 5’UTRs of *D. melanogaster* and then determined their presence or absence in the corresponding positions in the other species based on maf. AUG triplets that overlapped with any annotated CDS region were removed. The transcriptome and translatome data was obtained from the previous study ^42^.

### Generation of uORF-KO strain

We searched for possible sgRNA target sites near the start codon of the uORFs of *Clk* and *cyc* genes using the Benchling website (https://www.benchling.com/crispr/) to design optimal single guide RNA (sgRNA) sequences with high specificity and low off-target effects. Two pairs of sgRNAs were used for each gene. We then synthesized single-stranded complementary DNAs (ssDNAs) and annealed them to obtain double-stranded DNA (dsDNA), which served as the template for sgRNA expression. The dsDNA was then ligated into the BbsI-digested pU6B vector. The template sequences of the sgRNA uATG-KO are listed in SI Appendix, Table S5. The screening procedures of mutant flies are provided in the (SI Appendix, SI Methods).

### uORF mutation sequencing validation

The mutant fly was homogenized with DNA sequencing extract buffer [10 mM Tris-HCl at pH 8.0, 1 mM EDTA at pH 8.0, 25 mM NaCl and 0.02 % proteinase K (CW2584, CWBIO)] and incubated at 37°C for 30 min. PCR was performed with Taq Plus MasterMix (CW2849, CWbiotech). The PCR reaction was performed as follows: 94 °C for 2 min followed by 94 °C for 10 s, 57°C for 15 s, and 72°C for 1 min 20 s for 35 cycles. The primers used were listed in (SI Appendix, Table S5). PCR products were sequenced using commercial Sanger sequencing (AuGCT Co.).

### Cell culture and transfection

S2 cells were cultured in Schneider’s *Drosophila* Medium (Sigma) containing 10% heat-inactivated fetal bovine serum (FBS), 100 U/ml penicillin, and 100 µg/ml streptomycin (Thermo Fisher) at 25°C for 24 h to reach 2-4 × 10^6^ cells/ml before further treatments. Plasmid transfection was conducted with Lipofectamine 3000 (L3000001, Thermo Fisher) according to the supplier’s protocol.

### Dual-luciferase assay

The wild-type (WT) 5’UTR of *Clk* and *cyc* was cloned from cDNA by PCR, and uATG mutations were introduced into 5’UTR by the amplification primers. The PCR primers used for this purpose are listed in (SI Appendix, Table S5). The WT and mutated 5’UTR sequences were ligated into a linearized reporter plasmid (psiCHECK-2 vector, Promega). The entire sequences of all the plasmids were validated by Sanger sequencing. The *Renilla* luciferase activity associated with WT or uORF-mutated 5’UTRs was measured according to the manual of the Dual-Luciferase Reporter Assay System (Promega) 32 h after transfection and was normalized to the activity of firefly luciferase.

### Fly head collection

Flies were collected within 3-7 days of eclosion and entrained in LD for three days. After that, they were transferred into constant darkness (DD). Flies were collected during LD or on the first day of DD and frozen at −80°C. Frozen flies were vortexed for 10 s to separate the head from the body.

### Ribosome fraction analysis

The heads obtained from WT and mutant flies were collected and homogenized with lysis buffer, the lysates were clarified by centrifugation at 4°C and 20,000×g for 10 min, and the supernatants were applied to sucrose gradient ultracentrifugation. For each sample, the ratio of *Clk* and *cyc* mRNA abundance in the polysome fraction to that in the monosome fraction was calculated as the P-to-M ratio. Six biological replicates were performed for each sample. Details of the procedures can be found in the (SI Appendix, SI Methods).

### Western blot

Proteins were extracted from fly heads and cultured cells using SDS lysis buffer [50 mM Tris-HCl at pH 7.6, 150 mM NaCl, 1% Triton X-100, 0.5% SDS, 1 mM EDTA at pH8.0, 2 mM DTT, 1 × proteinase inhibitor and phosphatase inhibitor (4693116001 and 04906845001, Roche)]. After homogenization, protein lysates were centrifuged at 12,000 × *g* for 15 min at 4°C and incubated at 95°C in the loading buffer for 5 min. Equal amounts of protein were loaded into each well on 6% SDS-PAGE gels and then transferred to nitrocellulose membranes for 2 h at 90 V. Membranes were incubated with primary antibody at 4°C overnight, followed by secondary antibody at room temperature for 1 h. The primary antibodies used were as follows: guinea pig anti-CLK (1:1000, gift from Dr. Joanna Chiu) and rabbit anti-ACTIN 57B (1:5000, AC026, ABclonal, CN). Donkey secondary antibodies (1:10000 dilution) were conjugated either with IRDye 680 or IRDye 800 (926-68072 and 926-32213, LI-COR Biosciences, US). Bands were visualized and quantified with an Odyssey Infrared Imaging System (LI-COR Biosciences).

### Model simulation for the outcome of enhanced *Clk* translation

We adopted a previously published mathematical model of the *Drosophila* circadian clock ^47^. This system relies on a nonlinear, autonomous, first-order system of ordinary differential equations to simulate the regulatory weights among mRNAs or protein molecules in the molecular clock network. Details of the simulation can be found in the (SI Appendix, SI Methods).

### Total RNA extraction and RT-qPCR

RNA was extracted from frozen fly heads maintained at −80°C. 50 fly heads per sample were homogenized in Total RNA Isolation (TRIzol) Reagent (15596026, Invitrogen) by using a homogenizer with plastic beads. After mixing with trichloromethane, homogenates were centrifuged at 12,000 × *g* and the suspension was precipitated with 75% ethanol. After air dry, total RNA was reverse-transcribed into cDNA and genomic DNA was removed with One-step cDNA Synthesis SuperMix (AT311, Transgen). Quantitative real-time PCR was performed with One-Step RT-PCR SuperMix (AT141, Transgen). The PCR reaction was performed as follows: 45°C for 5 min; 94°C for 2 min; 94°C for 5 s, 60°C for 15 s, 72°C for 20 s for 40 cycles (Applied Biosystems). The ΔΔCT method was used for quantification. *Beta-Actin* was used as an internal control. The primers used are listed in (SI Appendix, Table S5).

### Locomotor activity monitoring and analysis

Locomotor activity levels of adult flies were monitored by *Drosophila* Activity Monitoring System (TriKinetics) for 7 days of LD followed by 7 days of DD. For DD rhythmicity, chi-squared periodogram analyses were performed by Clocklab (Actimetrics). Rhythmic flies were defined as those in which the chi-squared power was ≥ 10 above the significance line. Period calculations considered only rhythmic flies. Dead flies were defined by 0 activity on DD7 and removed from the analysis.

### Sleep measurement and analysis

*Drosophila* Activity Monitor system (Trikinetics) was used to analyze fly sleep. Virgin female and male flies of 3∼5 days old were used for the experiments unless otherwise specified. Flies were entrained under LD at 25°C for 3 days, and then their activities in the next 3 days under LD conditions were analyzed.

For sleep analysis during starvation, 3∼5 days old virgin females were used. After 3 days of entrainment in monitoring tubes with standard media (2% agar and 5% sucrose), baseline sleep was measured for 2 days. Standard food was then replaced at ZT0 by starvation media (2% agar) for 24 h. Flies were then re-fed with standard media for an additional 24-hour recovery period.

Sleep is defined as 5 min consecutive inactivity. Sleep was analyzed with SleepMat following previously published protocol ^60^.

### RNA-seq and circadian analysis

We collected heads from *Clk-*uORF-KO mutants and WT flies raised under LD cycles at 25°C at ZT 0, 4, 8, 12, 16 and 20. Library construction and sequencing with PE150 were conducted by Annoroad on the Illumina Nova6000 platform. Three biological replicates were sequenced for each sample. The clean data were mapped to the reference genome of *D. melanogaster* (FlyBase, r6.04) using STAR ^61^. Reads mapped to the exons of each gene were tabulated with htseq-count ^62^. Rhythmically expressed genes were identified by JTK_CYCLE under the threshold of < 0.05 on the permutation-based *P* values of the Jonckheere-Terpstra test (ADJ.P) implemented in the JTK CYCLE.

### GO-based enrichment

For GO-based enrichment analysis, GO annotation files were downloaded on 15 November 2023 from the Gene Ontology Resource (http://geneontology.org/)^63^ and contained 14,736 fly genes with at least one GO term. For each GO term *t*, we defined the following:

*N* = the total number of genes in the background distribution.

*n* = the total number of genes for GO analysis.

*M* = the total number of genes annotated by GO term *t*.

*m* = the number of genes annotated by GO term *t* in *n*.

The enrichment ratio (E-ratio) of *t* was then computed, and the *P* was calculated with the hypergeometric distribution as follows:

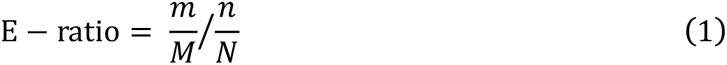

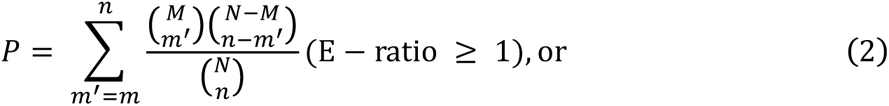

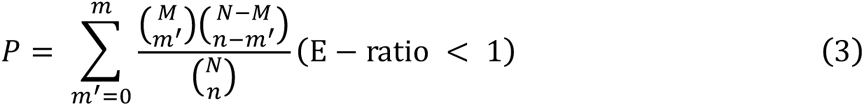

### Drug treatment

For pharmacological experiments, 5 mg/ml L-DOPA (BBI Life Sciences) was mixed in the fly food, and the flies were fed with L-Dopa food during sleep monitoring.

### Egg numbers quantification

Newly hatched virgins were picked out and allowed to mature for two days in separate vials. They were then mated by placing one virgin female with three male flies for two days. Subsequently, ten female parents from each group were transferred to a fresh dish containing a grape juice-based medium overlaid with yeast. This procedure was replicated in five separate dishes for each strain.

After one day of egg laying, the number of eggs deposited was meticulously documented. Following this assessment, the female parents were relocated to a new dish. This egg quantification process was conducted over a total duration of eight days. For the virgin female, egg counts were performed every two days, extending over a span of ten days.

### Quantification of offspring number per female fly

Newly hatched virgins were picked out and allowed to mature for two days in individual vials. They were then mated by placing one virgin female with three male flies for two days. After that, each female parent was transferred to a new vial and the number of offspring produced in 10 days was measured at 25°C and 29°C. All assays were performed with 20 females per genotype.

### Measurement of starvation resistance in adult flies

We selected 3-to 5-day-old adult males and females and placed them in a starvation medium (1.5% agar), with 10 flies per vial and 10 vials for both males and females from each strain. We observed every 6 hours or 12 hours to count the number of deaths under starvation conditions until all flies starved to death. The survival curves were plotted by the ggsurvplot package in R.

### Quantification and statistical analysis

Quantifications in all data graphs represent the mean of at least four biological replicates, and error bars represent the standard error of the mean (SEM). Two-tailed Mann-Whitney *U* test, Wilcoxon sign rank test, log-rank test, and two-tailed Student’s *t* test were used to calculate significance. All statistical analysis was carried out using *Statskingdom*. In each statistical analysis, the number of biological replicates and significant *P* values were noted in corresponding Fig. legends.

### Materials availability

All fly lines generated in this study are available from the Lead Contact.

