## Supplemental Information for "Upstream open reading frames dynamically modulate CLOCK protein translation to regulate circadian rhythm and sleep"

**This file includes:**

Supporting materials and methods
Figures S1 to S11
SI References

### Supporting Materials and Methods

#### Generation of uORF-KO strain

To achieve single base editing (ATG to ATT) by CRISPR/Cas9-mediated Homology-Directed Repair (HDR), a plasmid DNA donor with about 2 kilobases flanking the targeted site was constructed following a previous protocol [1]. The donors and pU6B-sgRNA plasmid were co-injected into the embryos of transgenic Cas9 flies collected within one hour of laying at the TsingHua Fly Center as described in Ni *et al.* [2]. The injected embryos were kept at 25°C and 60% humidity until adulthood (G0). The G0 adult flies that hatched from the injected embryos were individually crossed with another strain (*y sc v*) to increase the number of offspring. Then, the F1 progeny were crossed with flies carrying an appropriate balancer (*Dr, e/TM3, Sb*). After F2 spawning, the F1 individuals were screened for mutations of interest by genotyping. The primers used for genotyping are listed in the Key resources table. The F2 progeny whose parents showed positive genotyping results were then screened for the $y^-$ allele to separate the chromosome carrying *nos-Cas9*. The screened F2 males were crossed with flies containing the same balancer as mentioned above. After F2 genotyping, the progeny (F3) of positive F2 individuals carrying the same mutation were crossed individually to generate homozygous mutants in the F4 generation. The original homozygous mutants were backcrossed with Canton S flies for 9 generations.

#### Ribosome fraction analysis

The heads obtained from WT and mutant flies were collected and homogenized in a Dounce homogenizer with lysis buffer [50 mM Tris pH 7.5, 150 mM NaCl, 5 mM $MgCl_2$, 1% Triton X-100, 2 mM dithiothreitol (DTT), 20 U/ml SuperaseIn (Ambion), 0.5 tablets of proteinase inhibitor (Roche), 100 μg/ml emetine (Sigma Aldrich), and 50 μM guanosine 5′-[β,γ-imido]triphosphate trisodium salt hydrate (GMP-PNP) (Sigma Aldrich)] at 4°C. The lysates were clarified by centrifugation at 4°C and 20,000×g for 10 min, and the supernatants were transferred to new 1.5 ml tubes. 10-45% sucrose gradients were prepared in buffer (250 mM NaCl, 50 mM Tris pH 7.5, 15 mM $MgCl_2$, 0.5 mM DTT, 12 U/ml RNaseOUT, 0.5 tablets of protease inhibitor, and 20 μg/ml emetine) using a Gradient Master (Biocomp Instruments) in ULTRA-CLEAR Thinwall Tubes (Beckman Coulter). A sample volume of up to 500 μl was applied to the top of each gradient. After ultracentrifugation with a Hitachi P40ST rotor at 35,000 × rpm for 3 h at 4°C, the monosome and polysome fractions were collected, flash-frozen in liquid nitrogen, and stored at -80°C until further use.

The RNA in the monosome and polysome fractions was extracted separately using TRIzol reagent (Life Technologies, Inc.) and chloroform (Beijing Chemical Works) following the manufacturer's instructions and were reverse transcribed into cDNA using the PrimeScript™ II 1st Strand cDNA Synthesis Kit (Takara). RT-qPCR analysis of *Clk* and *cyc* cDNA was performed using PowerUp™ SYBR™ Green Master Mix (Thermo Fisher) following the manufacturer's instructions. The primer sequences employed for RT-qPCR are listed in Key resources table. For each sample, the ratio of *Clk* and *cyc* mRNA abundance in the polysome fraction to that in the monosome fraction was calculated as the P-to-M ratio. Six biological replicates were performed for each sample.

**Model simulation for the outcome of enhanced *Clk* translation**

Briefly, assuming that genes or proteins $j \in \{1,\ldots, n\}$ regulate the generation of gene/protein *i*, the subsequent type of differential equations as a general model were employed:

$$\frac{dx_i(t)}{dx} = \rho_i g\left(\sum_{j=1}^{n} \lambda_{ji}\, x_j(t) - \delta_i x_i(t)\right) x_i(t)\big(s_i - x_i(t)\big), 1 \le j \le n, \quad (1)$$

where the state vector $x_i$ represents the concentration of molecule $i$ at its site of action. The parameters $\lambda_{ji}$ represent regulatory weights that indicate the influence of molecule *j* on the production rate of molecule *i*. Positive and negative values of $\lambda_{ji}$ indicate activating or repressing effects, respectively. The larger absolute value of $\lambda_{ji}$, the stronger effect. It is worth noting that $\lambda_{Clk \to \mathrm{CLK}}$ in the model is applicable to the regulation of *Clk* mRNA to CLK protein (i.e., translation efficiency) in our study. To mimic the enhanced CLK translation caused by uORFs removal, we increased the original $\lambda_{Clk \to \mathrm{CLK}}$ from 0.026 (baseline) to three larger arbitrary values: 0.027, 0.028 and 0.030, respectively.

An odd sigmoid function modulates the cumulative regulatory influences, g: ℝ→ℝ, of the form

$$g(\mu) = \frac{\mu}{\sqrt{1+\mu^2}} = tanh\left(ln\,(\mu + \sqrt{1+\mu^2})\right),$$

together with a parameter $\rho_i > 0$ that indicates the maximum rate of $i$ production. The model incorporates logistic terms $x_i(s_i - x_i)$, where constants $s_i \ge 0$ indicate the saturation level of molecule *i*. The real parameter $\delta_i$ is the decay rate of *i*.

Simulations were performed on MATLAB (The MathWorks, Natick, MA). The system of ordinary differential equations is solved numerically by the subroutine ''ode45''. The actual equations and parameters are the same as those provided in the original model [3]. During each simulation cycle, the mRNA and protein levels of each molecule in the circadian network were recorded. We calculated the time intervals between the peaks of *tim* mRNA, serving as the proxy of circadian period length under the different regulatory weights.

### Supplementary figures

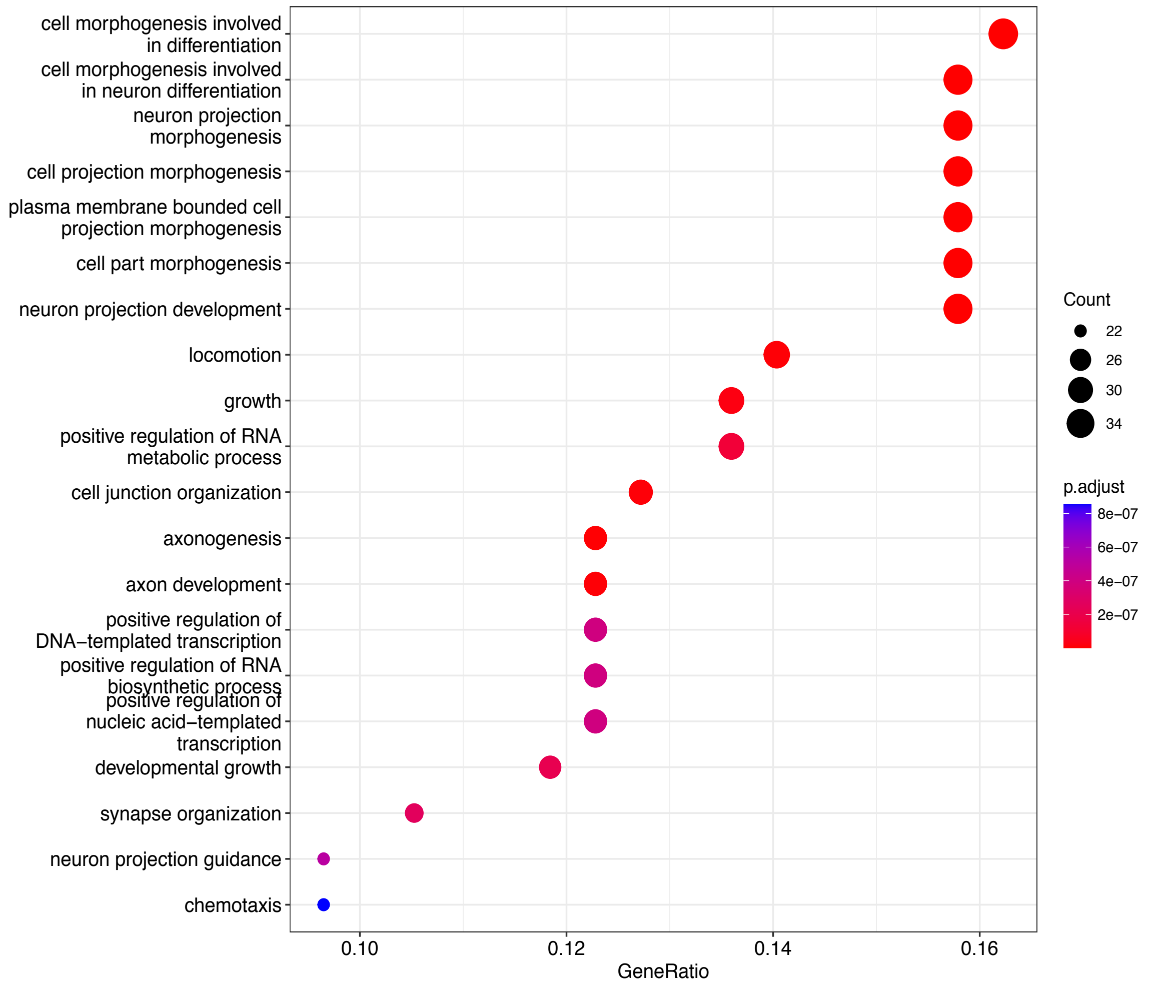


**Fig. S1. GO analysis for genes containing conserved uORFs.**

GO-based enrichment analysis for genes containing uORFs conserved across 23 *Drosophila* species.

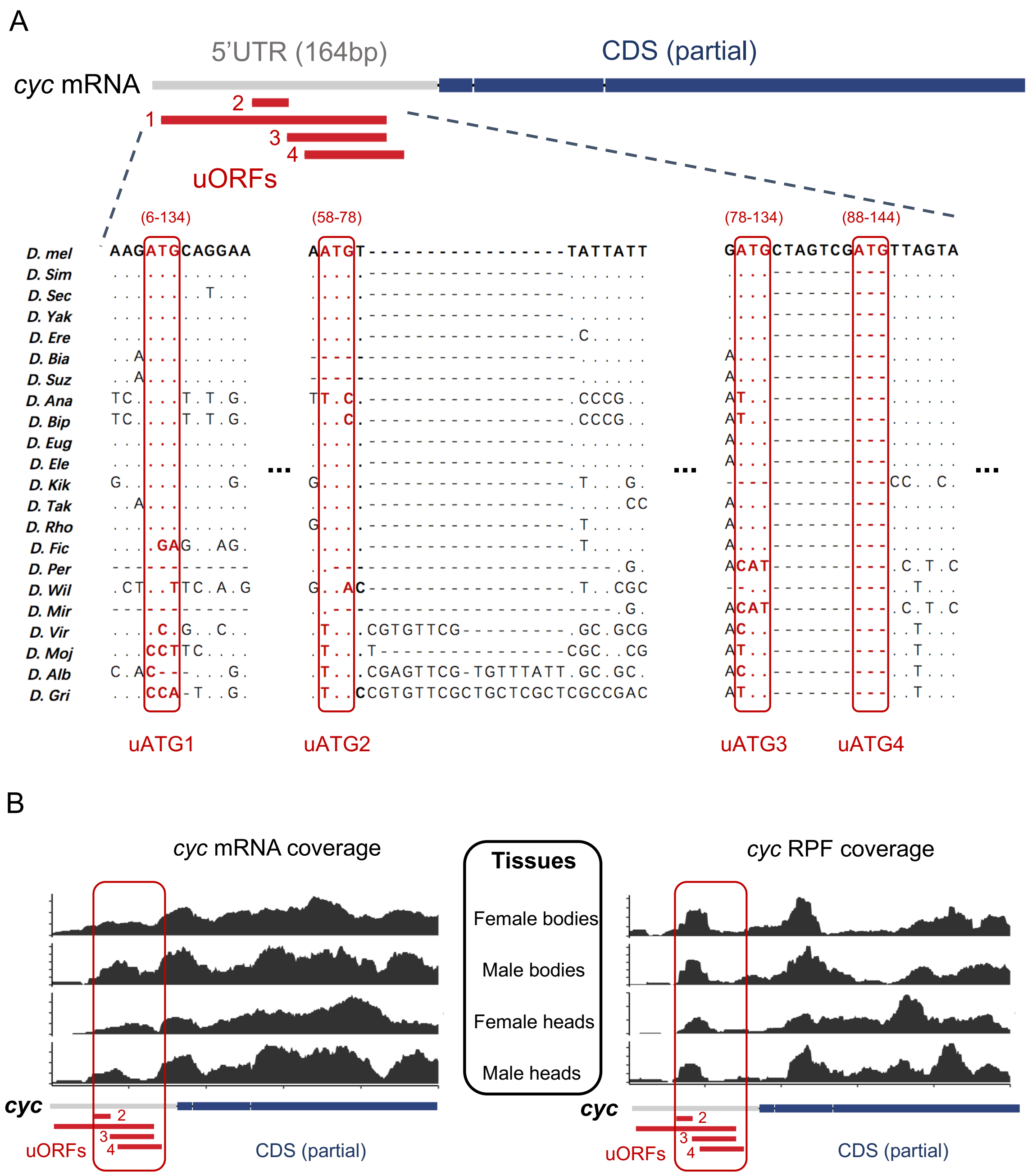


**Fig. S2. The conservation and translation of *cyc* uORFs in *D. melanogaster.***

(A) MSA of *cyc* uORFs among 23 *Drosophila* species. The start codons (uATGs) of uORFs are highlighted by red boxes. The position schemes of uORFs and partial CDSs are denoted above the MSA, with red and blue colors, respectively. The start and ending positions of each uORF (separated by a “-”) are given in the parenthesis above each uATG.

(B) The mRNA reads coverage (left) and ribosome-protected footprints (RPF) coverage (right) of uORFs of *cyc* mRNA from the heads and bodies. The position schemes of uORFs and partial CDSs are denoted at the bottom.

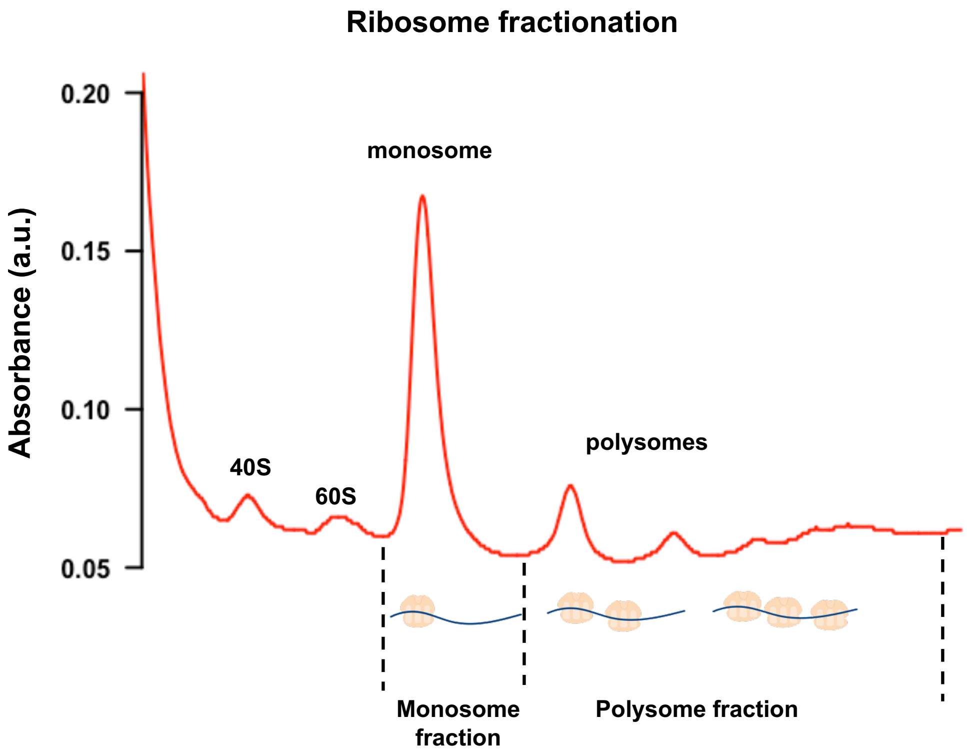


$$\text{P-to-M ratio} = \frac{\text{mRNA abundance in polysome fraction}}{\text{mRNA abundance in monosome fraction}}$$

**Fig. S3. Diagram demonstrating the separation of monosomes and polysomes by ribosome fractionation.**

Diagram illustrating the separation of monosomes and polysomes in a sucrose density gradient (10% - 45%). The P-to-M ratio is the ratio of mRNA abundance in the polysome fraction to that in the monosome fraction.

A

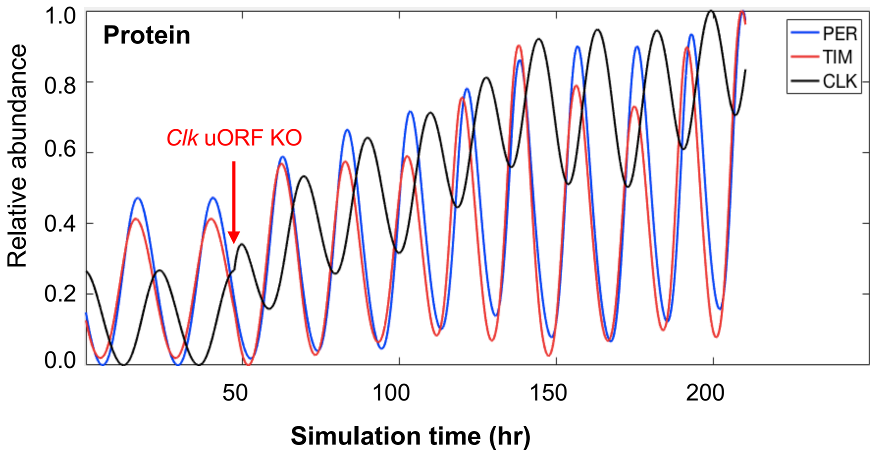


B

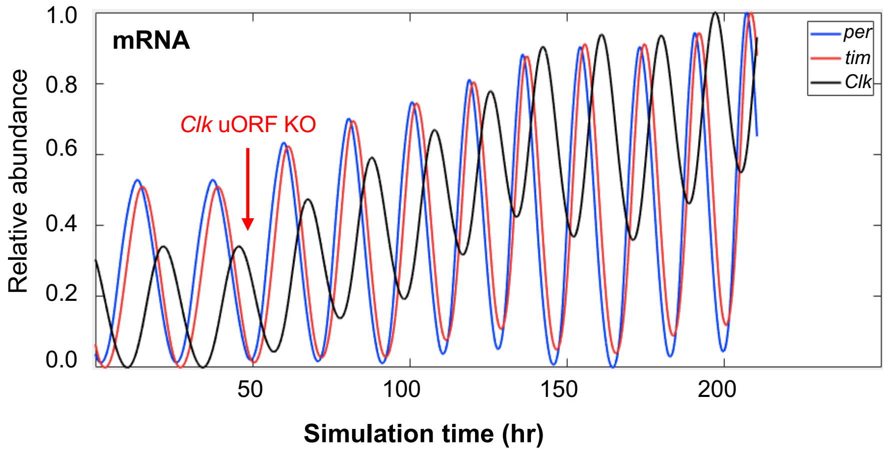


**Fig. S4. The simulated mRNA and protein abundance of *Clk*, *tim*, and *per* upon *Clk* uORF KO.**

(A) The simulated relative protein abundance levels of CLK, PER, and TIM along simulation time (hr) upon *Clk* uORF KO (see Fig. 4A for model details).

(B) The simulated relative mRNA abundance levels of *Clk*, *per*, and *tim* along simulation time (hr) upon *Clk* uORF KO (see Fig. 4A for model details).

The time point of increasing regulatory weight from *Clk* mRNA to CLK protein is marked by the red arrow labeled with *Clk* uORF KO.

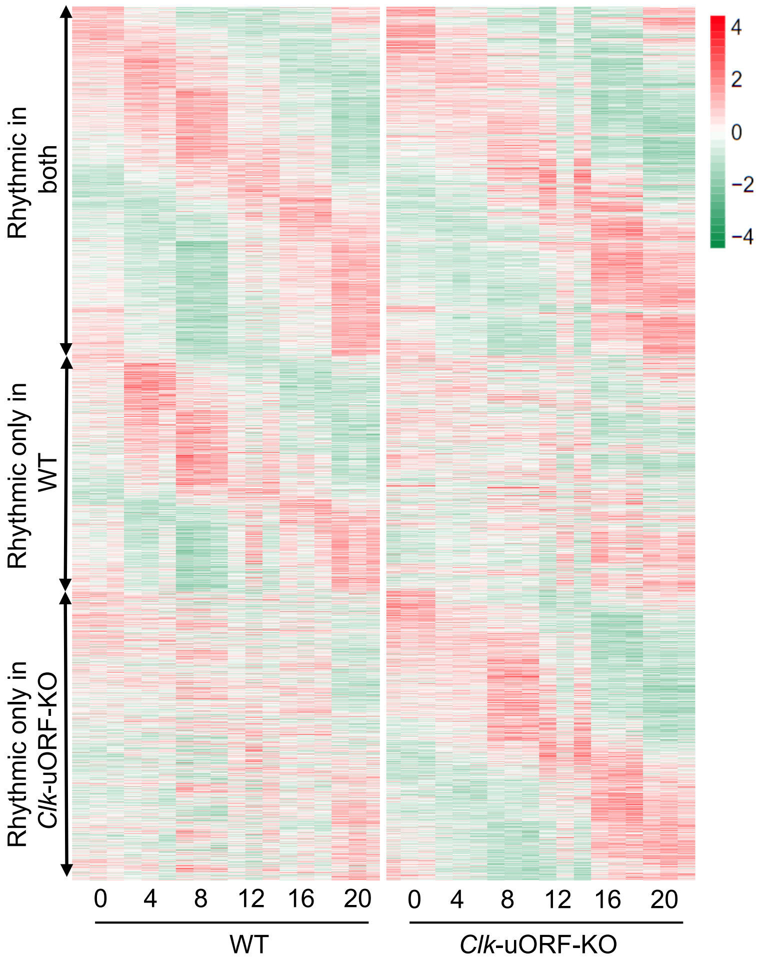


**Fig. S5. Rhythmically expressed genes in the heads of male WT and *Clk*-uORF-KO flies.**
A hierarchical clustering of the rhythmically expressed genes ordered by the phase of the oscillation. Values of genes are color-coded based on the intensities. Colors indicate low (green) and high (red) Z-scores.

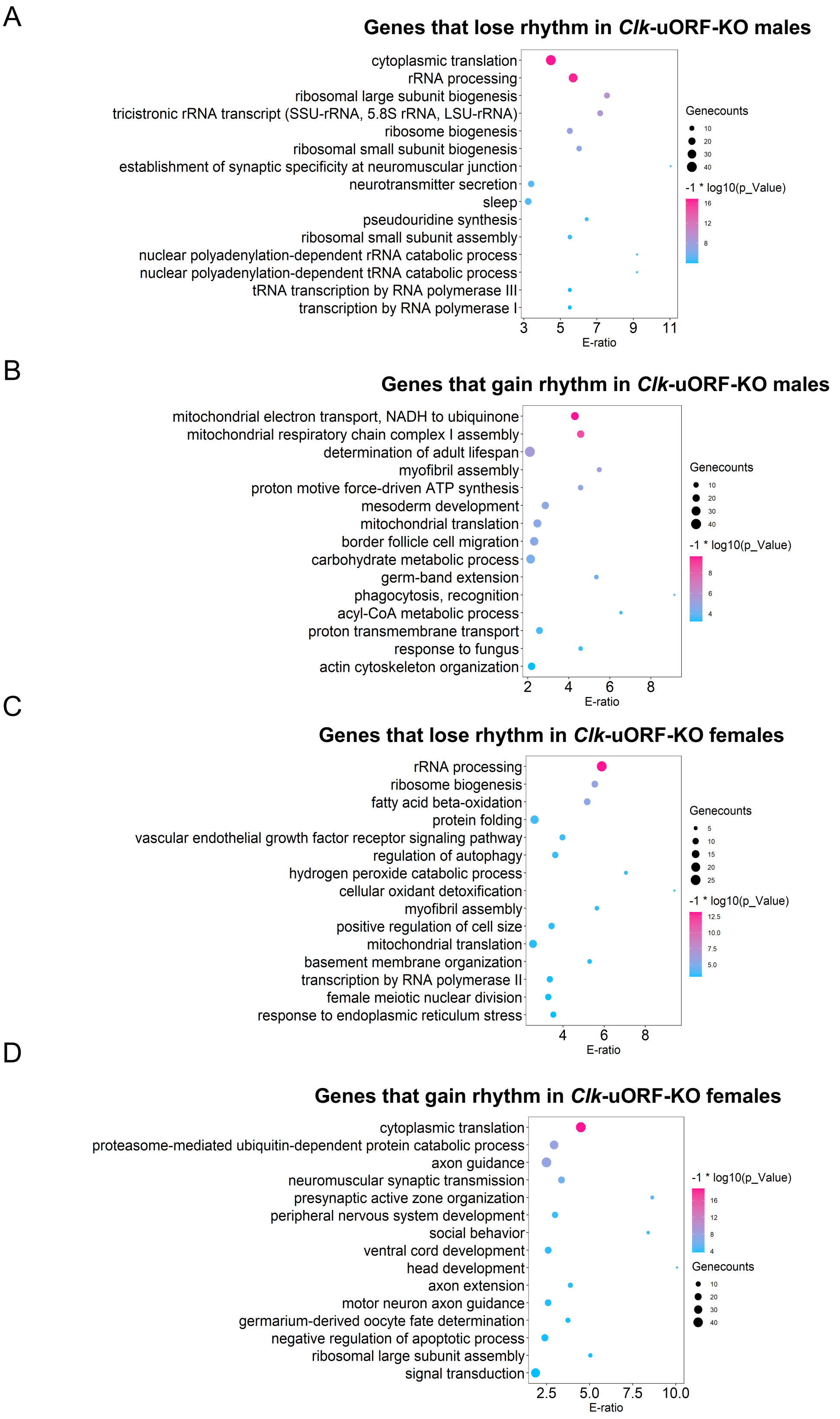
A
Genes that lose rhythm in *Clk*-uORF-KO males
cytoplasmic translation
rRNA processing
ribosomal large subunit biogenesis
tricistronic rRNA transcript (SSU-rRNA, 5.8S rRNA, LSU-rRNA)
ribosome biogenesis
ribosomal small subunit biogenesis
establishment of synaptic specificity at neuromuscular junction
neurotransmitter secretion
sleep
pseudouridine synthesis
ribosomal small subunit assembly
nuclear polyadenylation-dependent rRNA catabolic process
nuclear polyadenylation-dependent tRNA catabolic process
tRNA transcription by RNA polymerase III
transcription by RNA polymerase I
3
5
7
9
11
E-ratio
Genecounts
10
20
30
40
-1 * log10(p_Value)
16
12
8
B
Genes that gain rhythm in *Clk*-uORF-KO males
mitochondrial electron transport, NADH to ubiquinone
mitochondrial respiratory chain complex I assembly
determination of adult lifespan
myofibril assembly
proton motive force-driven ATP synthesis
mesoderm development
mitochondrial translation
border follicle cell migration
carbohydrate metabolic process
germ-band extension
phagocytosis, recognition
acyl-CoA metabolic process
proton transmembrane transport
response to fungus
actin cytoskeleton organization
2
4
6
8
E-ratio
Genecounts
10
20
30
40
-1 * log10(p_Value)
8
6
4
C
Genes that lose rhythm in *Clk*-uORF-KO females
rRNA processing
ribosome biogenesis
fatty acid beta-oxidation
protein folding
vascular endothelial growth factor receptor signaling pathway
regulation of autophagy
hydrogen peroxide catabolic process
cellular oxidant detoxification
myofibril assembly
positive regulation of cell size
mitochondrial translation
basement membrane organization
transcription by RNA polymerase II
female meiotic nuclear division
response to endoplasmic reticulum stress
4
6
8
E-ratio
Genecounts
5
10
15
20
25
-1 * log10(p_Value)
12.5
10.0
7.5
5.0
D
Genes that gain rhythm in *Clk*-uORF-KO females
cytoplasmic translation
proteasome-mediated ubiquitin-dependent protein catabolic process
axon guidance
neuromuscular synaptic transmission
presynaptic active zone organization
peripheral nervous system development
social behavior
ventral cord development
head development
axon extension
motor neuron axon guidance
germarium-derived oocyte fate determination
negative regulation of apoptotic process
ribosomal large subunit assembly
signal transduction
2.5
5.0
7.5
10.0
E-ratio
-1 * log10(p_Value)
16
12
8
4
Genecounts
10
20
30
40

**Fig. S6. The GO analysis for rhythmically expressed genes in WT and *Clk*-uORF-KO heads.**
(A-B) Gene Ontology (GO) enrichment analysis for genes that lose (A) or gain (B) rhythmic expression in the heads of male *Clk*-uORF-KO flies compared to WT.
(C-D) GO enrichment analysis for genes that lose (C) or gain (D) rhythmic expression in the heads of female *Clk*-uORF-KO flies compared to WT.

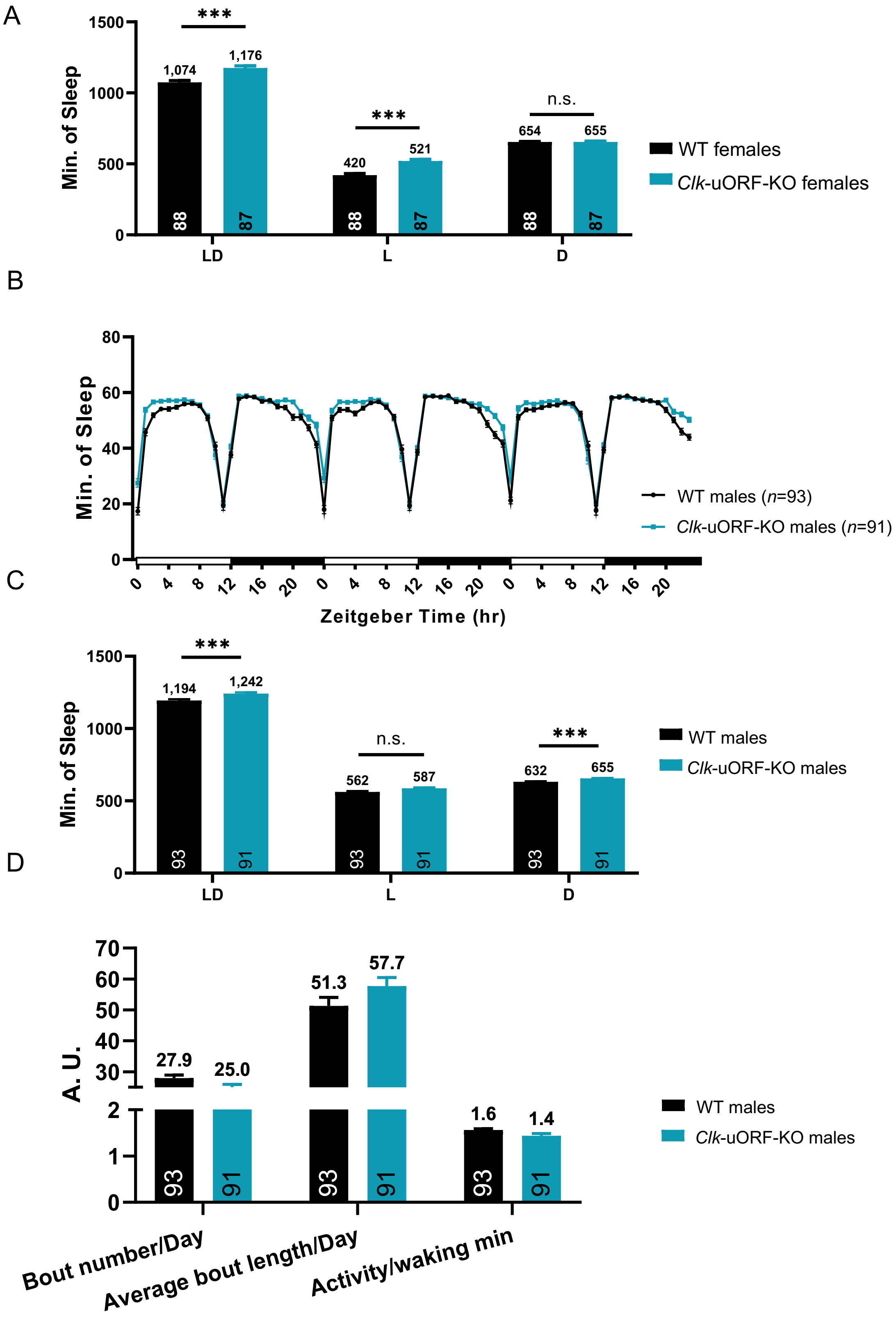


**Fig. S7. Knocking out *Clk* uORF does not substantially alter sleep in males.**

(A) Sleep duration during daytime (L), nighttime (D), and the entire day (LD) of female *Clk*-uORF-KO and WT flies. It matchs with Fig 6A and 6B.

(B) Sleep profile of male *Clk*-uORF-KO and WT flies under LD conditions.

(C) Sleep duration during daytime (L), nighttime (D), and the entire day (LD) of male *Clk*-uORF-KO and WT flies under.

(D) Daily sleep bout number, average bout length, and waking activity of male *Clk*-uORF-KO and WT flies under LD condition.

Data are presented as the mean ± SEM. The numbers of flies tested are shown in the brackets of legend or the bottom of histograms. Asterisks indicate statistical significance (two-tailed Student's *t* test. *, $P < 0.05$; ***, $P < 0.001$; n.s., $P > 0.05$).

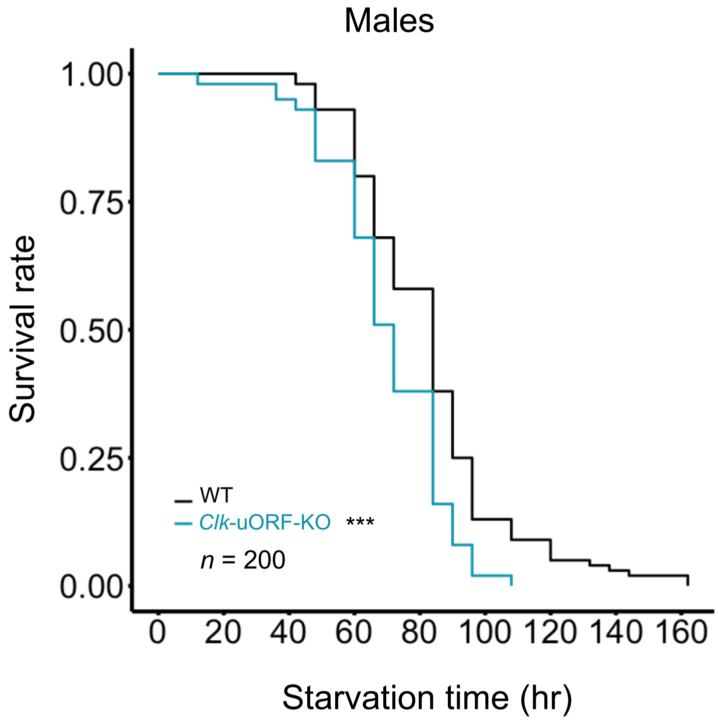


**Fig. S8. Knocking out *Clk* uORFs reduced starvation resistance in male flies.**

Survival curves of male flies under starvation conditions ($n = 200$; log-rank test; ***, $P < 0.001$).

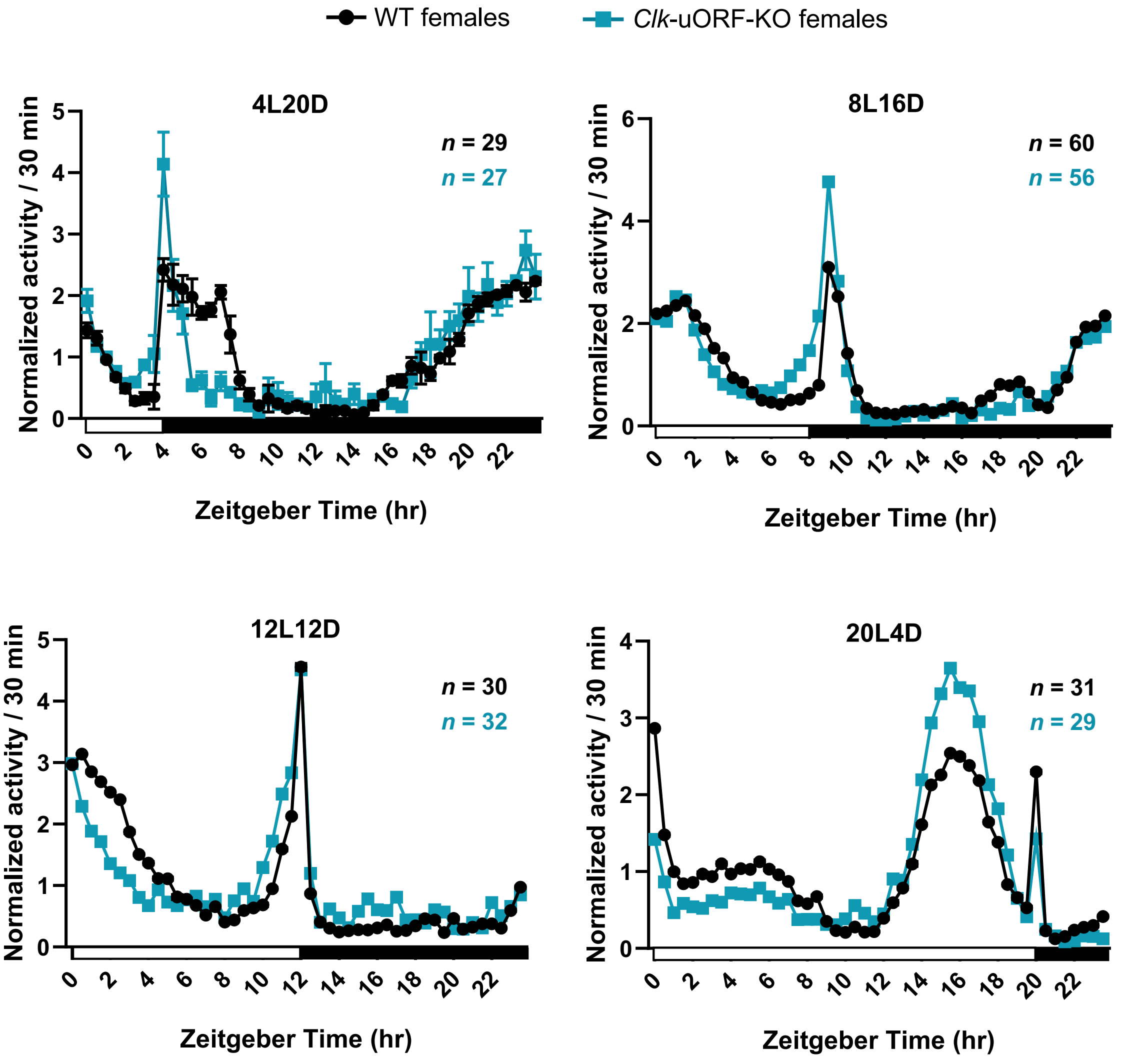


**Fig. S9. Knocking out *Clk* uORFs does not alter the adaptation of locomotor rhythm to seasonal photoperiod changes.**

Locomotor activity profiles of female WT and *Clk*-uORF-KO female flies under 4 h L : 20 h D (4L20D), 8 h L : 16 h D (8L16D), 12 h L : 12 h D (12L12D) and 20 h L : 4 h D (20L4D) conditions. The numbers of flies tested (*n*) are shown at the top-right of plots.

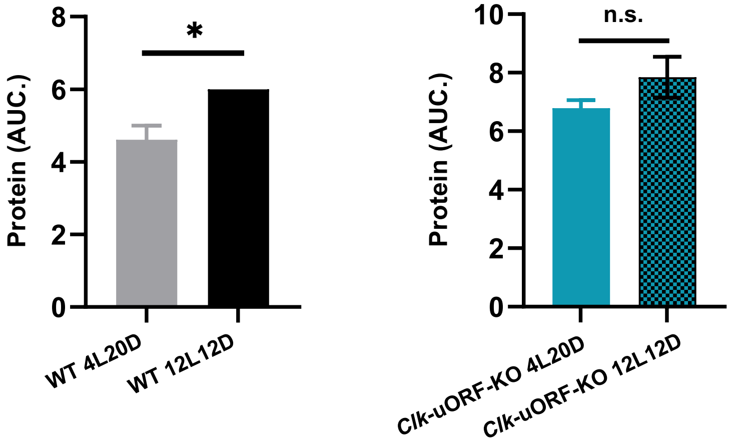


**Fig. S10. AUC analysis of daily CLK protein level under different photoperiods.**

AUC analysis of daily CLK protein level of female WT (left) and *Clk*-uORF-KO (right) flies under different photoperiods shown in Fig 8F. Each time point includes four biological replicates and data are presented as the mean ± SEM. Asterisks indicate statistical significance (two-tailed Mann-Whitney *U* test for unpaired comparisons. *, $P < 0.05$).

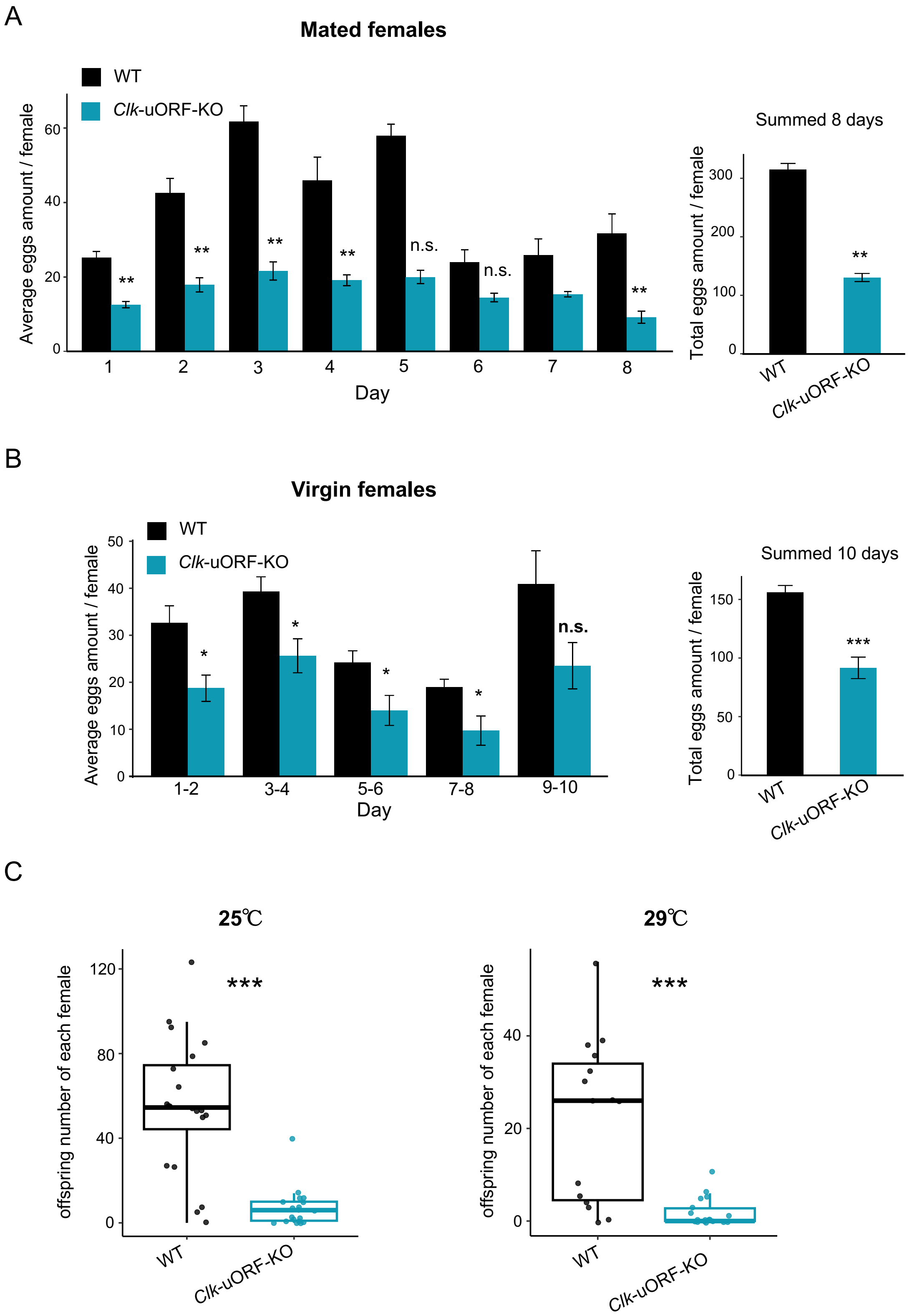
A
Mated females
WT
Clk-uORF-KO
Average eggs amount / female
0
20
40
60
**
**
**
**
n.s.
n.s.
**
1
2
3
4
5
6
7
8
Day
Summed 8 days
Total eggs amount / female
0
100
200
300
**
WT
Clk-uORF-KO
B
Virgin females
WT
Clk-uORF-KO
Average eggs amount / female
0
10
20
30
40
50
*
*
*
*
n.s.
1-2
3-4
5-6
7-8
9-10
Day
Summed 10 days
Total eggs amount / female
0
50
100
150
***
WT
Clk-uORF-KO
C
25℃
offspring number of each female
0
40
80
120
***
WT
Clk-uORF-KO
29℃
offspring number of each female
0
20
40
***
WT
Clk-uORF-KO

**Fig. S11. Knocking out *Clk* uORFs impairs fecundity.**

(A) The average 1-day egg number (left) and summed 8-day number (right) laid by a mated female of *Clk*-uORF-KO and WT over 8 consecutive days. Data are presented as the mean ± SEM ($n$=50; Wilcoxon rank sum test; **, $P < 0.01$; n.s., $P > 0.05$).

(B) The average 2-day egg number (left) and summed 10-day number (right) laid by a virgin of *Clk*-uORF-KO and WT over 10 consecutive days. Data are presented as the mean ± SEM ($n$=50; Wilcoxon rank sum test; *, $P < 0.05$; ***, $P < 0.001$; n.s., $P > 0.05$).

(C) The offspring number per female parent of *Clk*-uORF-KO and WT over 10 days at 25°C (left) and 29°C (right) ($n$=20; Wilcoxon rank sum test; ***, $P < 0.001$).
